# Homunculus Interruptus: A motor association area in the depth of the central sulcus

**DOI:** 10.1101/2022.11.20.517292

**Authors:** Michael A. Jensen, Harvey Huang, Gabriela Ojeda Valencia, Bryan T. Klassen, Max A. van den Boom, Timothy J. Kaufmann, Gerwin Schalk, Peter Brunner, Dora Hermes, Gregory A. Worrell, Kai J. Miller

## Abstract

Cells in the precentral gyrus of the human brain directly send signals to the periphery to generate movement and are topologically organized as a map of the body. We find that movement induced electrophysiological changes from implanted depth electrodes extend this map 3-dimensionally throughout the volume of the gyrus. Unexpectedly, this organization is interrupted by a motor association area in the depths of the central sulcus at its mid-lateral aspect that is active during many different types of movements from both sides of the body.

---

The organization of body movements on the convexity of the precentral gyrus (PCG), named the homunculus, was discovered nearly a century ago by direct brain surface stimulation in awake neurosurgical patients^1^, and follows a medial-to-lateral pattern of lower extremities, upper extremities, and face. Subsequent measurements of task driven changes in functional MRI imaging (fMRI), magnetoencephalography^2^, and brain surface electrophysiology with electrocorticography (ECoG) have all recapitulated this somatotopic organization^3–5^. Neurons from each of these somatotopic areas in the PCG, called primary motor cortex, communicate with the brainstem and spinal cord to produce body movements. *Primary* brain areas are generally defined by having a simple chain of synaptic connections to the peripheral body and a direct topographical mapping to the outside world: retinotopy in the calcarine cortex (visual), tonotopy in the transverse temporal gyrus (auditory), and somatotopy in the post-central gyrus (sensation) & the PCG (movement)^6^. Brain areas that can be related to these functions but are not themselves primary are called *association* areas. Association areas may or may not exhibit topographic organization and are often found to coordinate basic topographic features for a more complex purpose^7^. Our research began as an effort to simply characterize the PCG primary motor cortex electrophysiologically throughout its 3-dimensional volume, measuring from superficial and deep areas simultaneously. This was made possible by working with patients who have penetrating stereoelectroencephalographic (sEEG) depth electrodes placed in their brains for clinical practice. In these measurements, we expected to find only classic primary motor properties in the PCG, but instead found evidence for an association area interrupting the otherwise somatotopic representation.

In our treatment of patients with drug resistant focal epilepsy, sEEG depth electrodes may help to identify where in the brain seizures originate from and propagate to^8^. sEEG has emerged as the predominant implanted monitoring approach for epilepsy, largely replacing brain surface ECoG arrays in recent years^9^. The 0.8mm diameter sEEG electrodes are minimally traumatic, allow for volumetric characterization of seizure networks, and are well tolerated^10,11^. As a complement to electrical stimulation mapping, which perturbs the brain to characterize function, we also perform simple behavioral tasks from which we can analyze electrophysiological changes to map neural activity in the immediate vicinity of each electrode. The electrical potential signals measured by sEEG from cortex during behavior show the same general features as those seen in ECoG^12,13^: event-locked raw voltage deflections, oscillations (rhythms), broadband (power-law) spectral changes (Fig 1). As in ECoG, we find that, in peri-central areas, simple movements produce 1) decrease in power in narrowband oscillations in the ~10-40Hz range and 2) broadband spectral increases above ~50Hz that we capture between 65-115 Hz. Such broadband changes have been shown to be a general correlate of neural population firing rate^14^.

**Figure 1.**
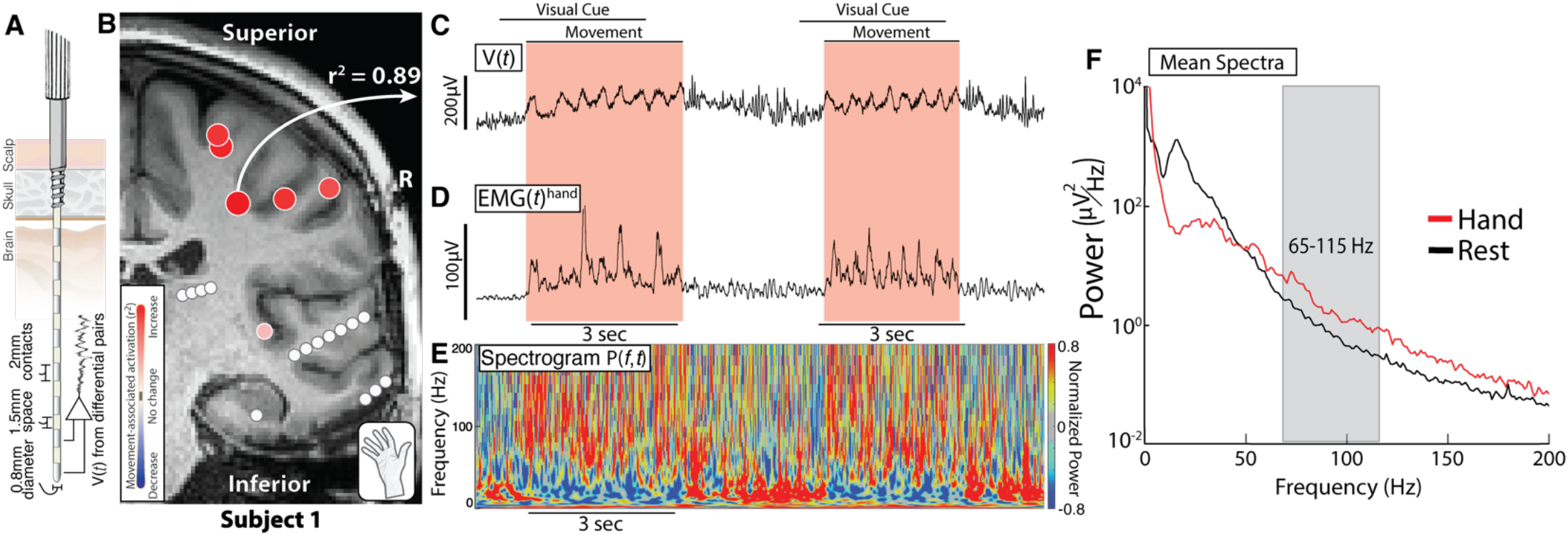
The stereoelectroencephalographic (sEEG) signal during movement. **A.** Schematic of an sEEG electrode demonstrating recording sites throughout the brain volume. **B.** Coronal T1 MRI cross section through the precentral gyrus and central sulcus with differential electrode pair channels (circles) showing active brain areas during hand movement (r^2^ values). **C.** Voltage timeseries measured from the channel in B. **D.** Rectified EMG signal measured from forearm extensors during simple hand movement. **E.** Spectrogram aligned to C and D shows increases in the power at high frequencies and decreases in power at low frequencies during movement. **F.** Power spectra density plot showing a broadband increase in power during movement at higher frequencies and a narrowband decrease in power during movement at lower frequencies. The gray box indicates the frequency range analyzed as a reflection of local brain activity.

**Figure 2.**
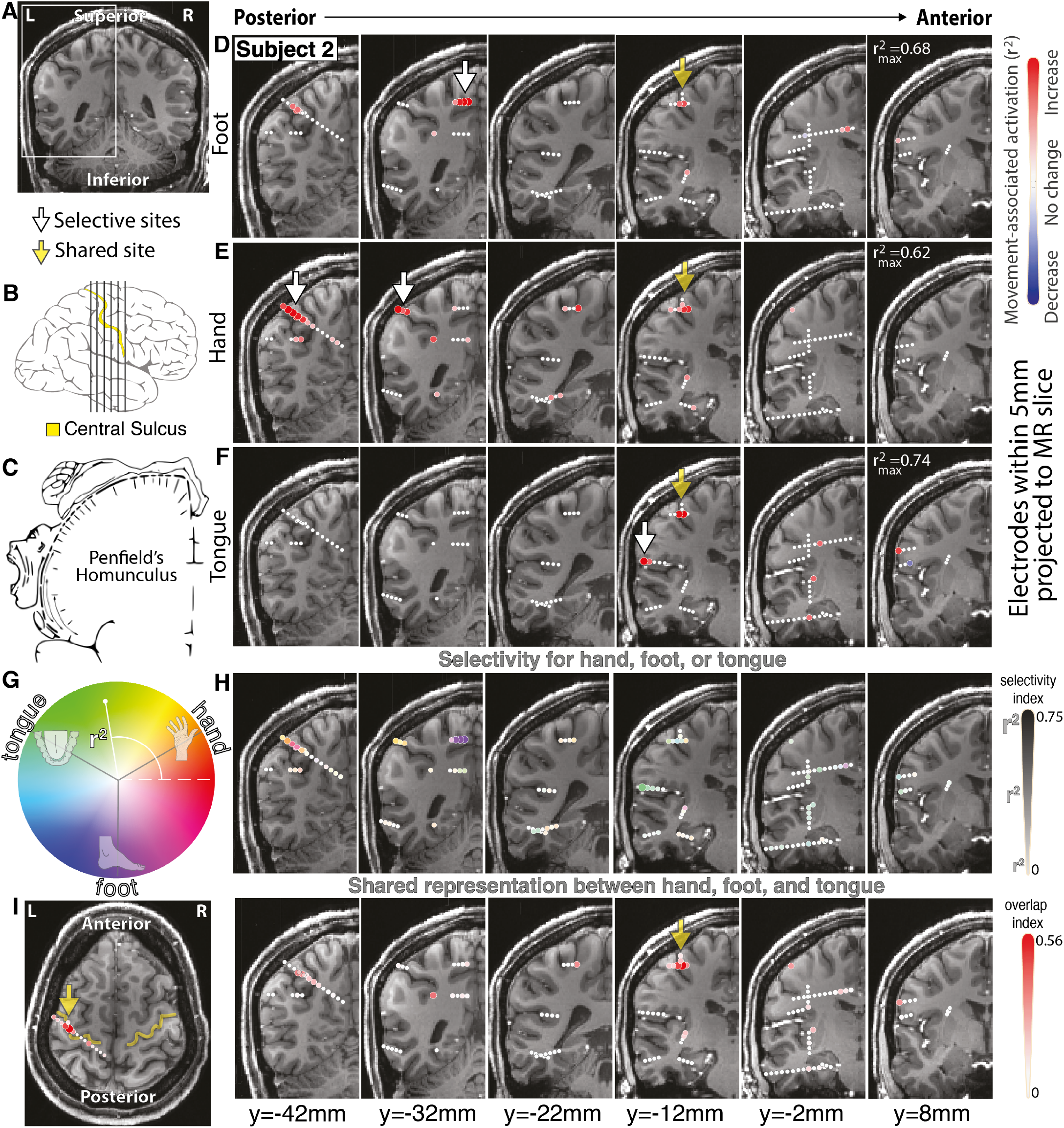
Volumetric electrophysiological changes during simple body movements. **A.** Coronal T1 MRI, with inset box assessed throughout figure. **B.** Coronal slices throughout central sulcus / precentral gyrus shown with black vertical lines. **C.** The classic Penfield motor homunculus from awake stimulation shown as reference^1^. **D-F.** Maps of sEEG power spectral change in the 65-115Hz range during foot, hand, and tongue movement, respectively. Maximum scaling of the colorbar is noted in the top right of each row. **G.** Circular colormap showing selectivity for hand, tongue or foot using projected r^2^ values into the complex plane for maps plotted in (H). Diameter and intensity indicate magnitude of selectivity, while color reflects selective movement type(s). Note that a channel that is equally active (even if highly so) during all 3 movement types will be plotted small and white, even if highly selective for each. **H.** Movement selectivity maps using scale from (G). The most selective sites from these plots are shown with white arrows in D-F **I.** Maps of shared activity in movement (geometric mean of hand, tongue, and foot r^2^ values). The peak of shared representation is shown with yellow arrows in D-F. Insignificant channels are plotted in white in all panels.

Subjects performed a simple block-designed task of randomly interleaved tongue, hand, or foot movements (contralateral to SEEG array) with rest in between while electromyography was recorded from each body area. Simple analysis of broadband changes extended the classic selective somatotopic representation of individual body parts into the sulcal depths, with foot along the midline, hand in the superior-lateral part, and tongue in the lateral aspect^1^ (Fig 2). In the depths of the central sulcus, at its mid-lateral aspect, the distinct topology of the homunculus was unexpectedly interrupted by a region that was electrophysiologically active during all three movement types. We call this the “Rolandic motor association” (RMA) area in reference to the historical name of the central sulcus (fissure of Rolando)^15^. This RMA area was clearly observed in all 13 participants (Fig 3) and was surrounded by movement-specific somatotopic regions in each case. RMA area is active during both ipsilateral and contralateral body movements (Fig 4, S8). While electrophysiological changes in classic somatotopic sites precede movement onset and peak early in movement blocks, RMA sites showed more nuanced dynamics throughout movement (Fig 4, S9). Standard clinical stimulation mapping incidentally included the RMA for one patient (Subject 3). In this patient, stimulation of the RMA did not disrupt movement or speech function (up to 5mA bipolar testing), though stimulation at primary motor foot-specific regions in the same patient produced muscle contraction at 2mA. This further indicates that the RMA is a motor association region rather than a primary motor site.

**Figure 3.**
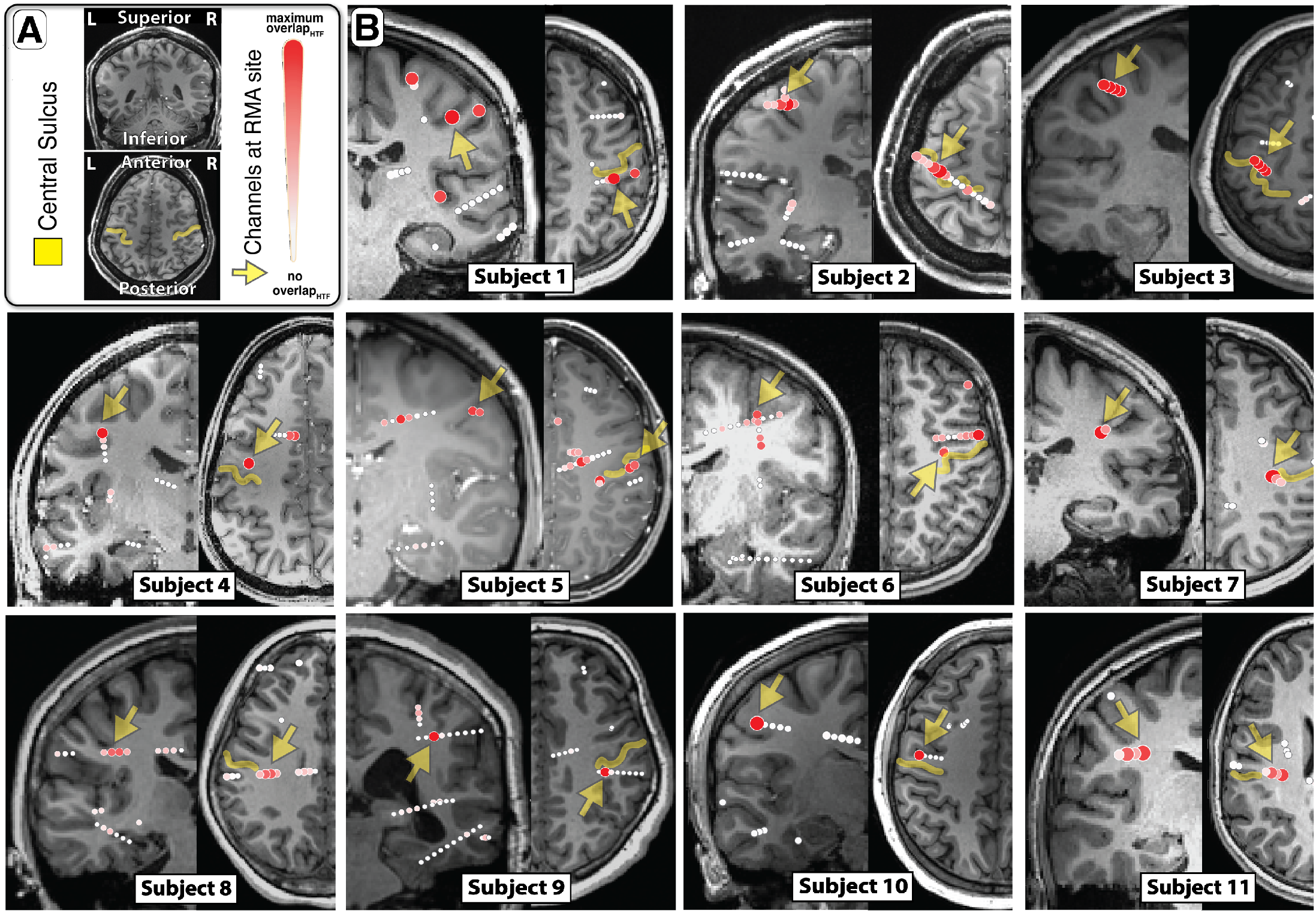
Rolandic motor association area within the central sulcus. **A.** Orientation of coronal and axial slices **B.** Axial and coronal views demonstrate localization of channels with shared representation between hand, tongue, and foot movement within the central sulcus. We call this region of shared representation the Rolandic motor association (RMA) area (yellow arrow).

**Figure 4.**
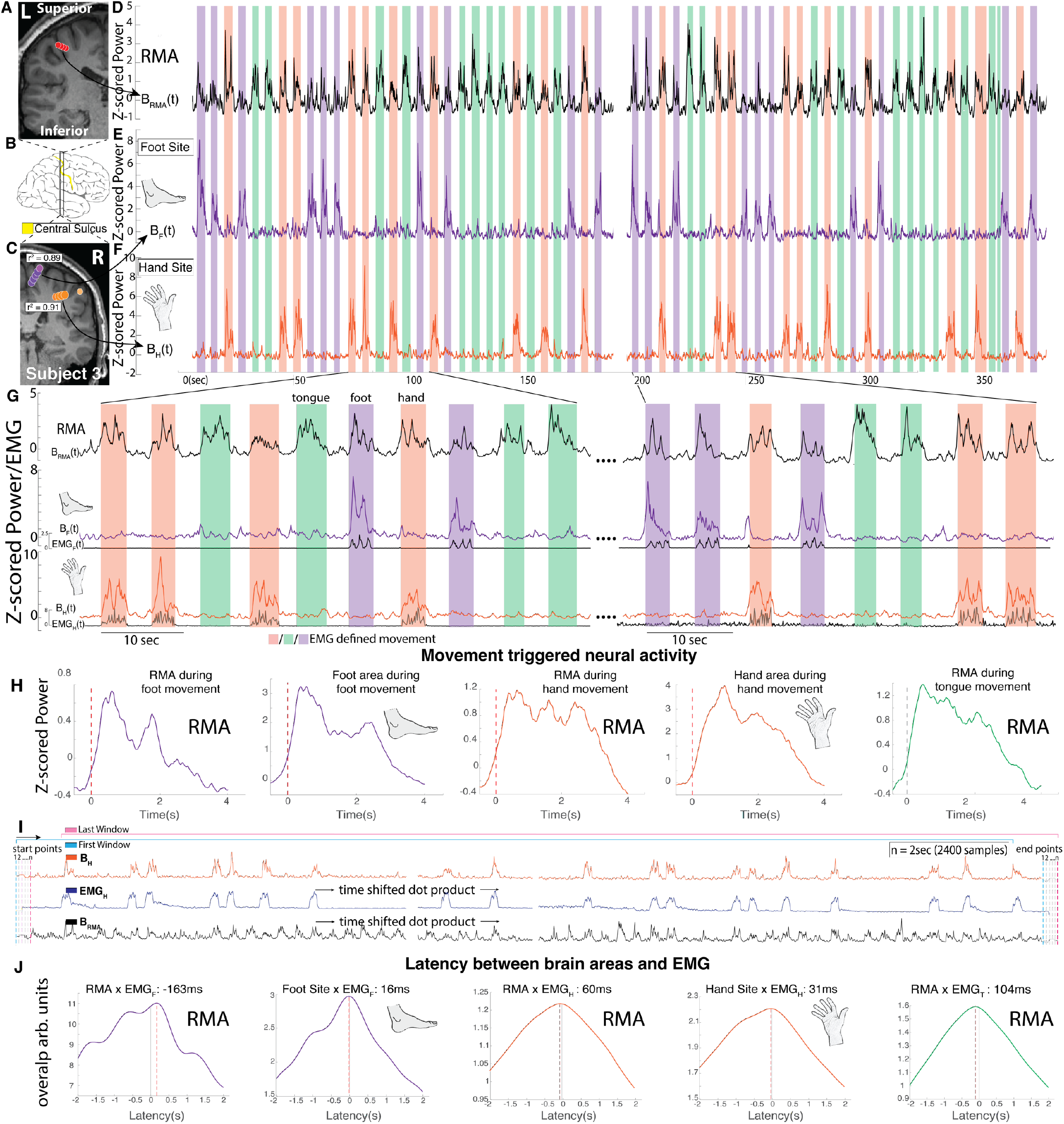
Temporal dynamics of neural activity in the precentral gyrus, subject 3. **A.** Coronal T1 MRI cross section through the precentral gyrus and central sulcus in the left hemisphere, with plotted shared activity revealing the RMA area, as in Fig 2H. **B.** Position of coronal slices in (A)&(C). **C.** Right hemisphere, with plotted selectivity revealing somatotopic hand and foot representation, as in Fig 2I. **D-F.** Timecourse of broadband activity (65-115Hz, “B”) for RMA, foot, and hand channels, reflecting local neural activity. Background shading indicates EMG-defined movement periods of the hand (red), foot (purple), and tongue (green). **G.** Magnified epochs from (D-F), with EMG included. **H.** Brain activity averaged to onset of foot, hand, and tongue EMG. **I.** Schematic of time-shifted dot products to calculate latencies between brain activity and movement. **J.** Profiles of latency between brain areas (RMA, foot selective, and hand selective) and EMG (hand and foot).

Our present measurements succinctly establish that RMA area is a somatotopically non-specific locus in an area traditionally thought to follow a clear map of distinct representation. Although many have suggested that the homunculus may not be as distinct as textbook illustrations^5^, arguments center around how blurred the overlap in the map is rather than a refutation of limb-specific representation^16^. Therefore, the lack of somatotopic separation in the RMA area is a fundamental exception to our organizational understanding of the human precentral gyrus and its relationship to movement.

This RMA area has likely been overlooked in prior studies because of accessibility of the cortical surface convexity and primary use of surface ECoG for awake motor mapping^1,17^. Nonetheless, two emerging studies may provide insight into the role of RMA area in motor circuitry. Silva et al. describe a region within the central sulcus that plays a premotor role in the coordination of speech production^18^. Our finding suggests that this may be a subfunction of the RMA area, though not the primary function, since the RMA area is active during foot and hand movement. Using a battery of tasks in fMRI, Gordon et al. propose that there are multiple precentral gyral regions interrupting the otherwise somatotopic representation, that coordinate whole-body action plans with specific connections to both striatal regions and the centromedian nucleus of the thalamus^19^. One of the PCG regions they suggest plays this role directly overlies the RMA area; another, more superiorly positioned region, is consistent with a secondary RMA region that we also observed in 5 subjects (Fig S10). Collectively, our RMA area finding, and the emerging work of others, clearly shows that the traditional view of the motor homunculus as an uninterrupted topography on the precentral gyrus must be revised. Future study to understand how the RMA plays a wider role in motor circuitry might begin with more nuanced experimental paradigms that explore how this region interacts with primary motor regions and other motor association areas during movement preparation^20^, motor imagery^21^, action observation^22^, and sensory feedback^23^.

## Funding

This work was supported by the Brain Research Foundation with a Fay/Frank Seed Grant (KJM), and by NIH-NCATS CTSA KL2 TR002379 (KJM), NIH P41-EB018783 (PB) & NIH U01-NS128612 (KJM, PB, GAW). Manuscript contents are solely the responsibility of the authors and do not necessarily represent the official views of the NIH. The funders had no role in study design, data collection and analysis, decision to publish, or preparation of the manuscript.

## Acknowledgments

We are grateful to the patients who volunteered their time to participate in this research, to Bambi Wessel, Cindy Nelson, and the staff at St. Marys hospital. The term “Homunculus Interruptus” was generously suggested by Jon Willie.

## MATERIALS and METHODS

### Ethics statement

The study was conducted according to the guidelines of the Declaration of Helsinki and approved by the Institutional Review Board of the Mayo Clinic IRB# 15-006530, which also authorizes sharing of the data. Each patient / representative voluntarily provided independent written informed consent to participate in this study as specifically described in the IRB review (with the consent form independently approved by the IRB).

### Subjects

Thirteen patients (6 females, 11-35 years of age - **Table S1**) participated in our study, each of whom underwent placement of 10-15 sEEG electrode leads for seizure network characterization in the treatment of drug resistant partial epilepsy. Electrode locations were planned by the clinical epilepsy team based on typical semiology, scalp EEG studies, and brain imaging. No plans were modified to accommodate research, nor were extra electrodes added. Thirteen of fifteen consecutive treated patients participated in our motor task. One excluded patient did not wish to participate in research (i.e. did not consent) and the other excluded patient did not have appropriate peri-central electrodes. All experiments were performed at the Mayo Clinic in Rochester, MN. Each patient or parental guardian provided informed consent as approved by the Institutional Review Board at Mayo Clinic (IRB# 15-006530). All T1 MRI sequences were defaced prior to uploading using an established technique^24^, to avoid potential identification of participants.

### Lead Placement, Electrode Localization, and Referencing

The platinum depth electrode contacts (DIXI Medical) were 0.8mm in diameter with 2mm length circumferential contacts separated by 1.5mm (**Fig 1, Fig S1**)^25^. Each lead contained 10-18 electrode contacts. Surgical targeting and implantations were performed in the standard clinical fashion^25^. Intraoperatively, anchoring bolts were placed stereotactically in 2.3 mm holes in the skull, and leads were then advanced to target through the bolts. Once at target, leads were secured into the skull by a guide screw and cap (**Fig 1, Fig S1**).

Electrode anatomic localizations were determined by co-registration of post-implant CT scan to pre-implant MRI. Each pre-operative T1 MRI was realigned to the anterior and posterior commissure stereotactic space (AC-PC) using VistaSoft^26^, then co-registered to the post-implant CT using SPM12^27^. The electrode positions in the merged MR-CT space were plotted in each figure using a custom open source MATLAB toolbox we developed (“SEEGVIEW”)^28^.

All data were re-referenced in a bipolar fashion, producing channels that reflect mixed activity between voltage timeseries measured at two adjacent electrode contact sites. Plotted points for brain activity in this study each represent an interpolated point between the two electrodes that make up each differential pair channel. Only adjacent differential pair channels were considered (i.e. 1.5mm from one another, on the same lead, and within the same lead segment for segmented leads) (**Fig S1, S5**). In each figure, channels were plotted using SEEGVIEW, which slices brain renderings, and projects channels to the center of the closest slice^28^. The purpose of this is to present analyses in an interpretable, clinically familiar manner. This projection approach imposes a longer projection distance if fewer/thicker slices are chosen for visualization. With fewer slices, all projected channels can be viewed more simply. Note that a channel reflecting activity in the gray matter at the depth of a sulcus may appear to be in white matter. Anatomic features (central sulcus, etc) and designations of each channel were carried out by a neuroradiologist (TJK).

### Motor Task

Data were collected during a motor task involving 1) opening and closing of the hand, 2) side-to-side movement of the tongue with mouth closed, and 3) alternating dorsi- and plantar flexion of the foot (contralateral to the hemisphere with peri-central sEEG electrode coverage). Patients were visually cued to perform simple self-paced (~1Hz) movements in response to images of a hand, tongue, or foot, and to remain still during interleaved rest periods (blank black screen). Twenty cues (trials) of each movement type were shuffled in random order and move & rest cues were 3s in duration (**Fig S2**). This task was chosen based upon prior work, which has produced clear results in recordings from the brain surface^3^. The BCI2000 software was used for stimulus presentation and data synchronization^29^, with stimuli presented on a 53 x 33 cm screen, 80-100 cm from the face (**Fig S2)**. If patients were not participating with the task, the experimental run would be stopped and re-run later.

### Electrical stimulation mapping

In subject 3, stimulation mapping was performed at the RMA site for clinical purposes. We found that bipolar stimulation up to 5mA at the RMA site did not produce a sensory response and did not interrupt or elicit movement. Bipolar clinical stimulation at 2mA produced contraction of the anterior tibialis at multiple foot selective sites. Stimulation mapping was not performed in other patients as it was not included in the research protocol – though the IRB does allow for use of clinical data, provided that it preserves the privacy of the patient.

### Electrophysiological Recordings

Intracranial sEEG signals were initially recorded relative to a clinician-selected reference in the white matter away from tissue with likely seizure or motor involvement. Voltage timeseries were recorded with the 256-channel g.HiAmp amplifier (gTec, Graz, Austria). Recordings were sampled at 1200Hz, with an anti-aliasing filter which dampened the signal by 3dB at 225Hz.

Electromyography (EMG) was measured from the forearm flexors/extensors (hand), base of chin (tongue), and anterior tibialis (foot) during the motor task (**Fig S3**). All sEEG and electromyographic (EMG) signals were measured in parallel, and delivered to both the clinical system and the research DC amplifier (g.HIAmp system, gTec). sEEG and EMG signals were synchronized with the visual stimuli using the BCI2000 software^29^.

### Signal Processing and Analysis (Figs S4&S7)

#### Trial-by-trial power spectral analysis (Fig S4)

All analyses were performed in MATLAB. Adjacent electrode contacts were first bipolar re-referenced to neighboring contacts on the same lead segment **(Fig S5)**. To determine the precise timing of movement onset and offset in response to a visual cue, EMG-timing based analyses were chosen for behavioral analysis rather than the timing of the visual movement cue **(Fig S6)**. EMG measuring tongue movement was lacking in subject 6. In this case, we defined tongue movement periods by shifting all visual tongue cue onsets/offsets based on the subject specific average delay between the onset/offset of cue and EMG activity for hand and foot movements. Within each movement trial, averaged power spectral densities (PSDs) were calculated from 1 to 300 Hz every 1 Hz using Welch’s averaged periodogram method with 1 second Hann windows to attenuate edge effects^30^ and 0.5 second overlap. The averaged PSD for each movement or rest trial was normalized to the global mean across all trials. We normalized the PSDs in this way since brain signals of this type generally follow a 1/f, power law, shape^31^, so that lower frequency features dominate if un-normalized. From each of these normalized single trial PSDs, averaged power in a broadband high frequency band (65-115 Hz) was calculated for subsequent analysis, as previously described^32^. This captures broadband activity above the known range of most oscillations and avoids ambient line noise at 60 and 120Hz.

For each bipolar re-referenced channel, we calculated separate signed r^2^ cross-correlation values (r^2^) of the mean spectra from 65-155 for each movement modality. Each channel’s r^2^ value was determined by comparing mean power spectra between movement trials (separately) and rest. To minimize the cross-effects of beta rebound, movement trials of each type were only compared with rest trials that followed that same movement type^33^:

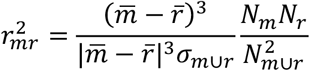

where *m* denotes power samples from movement, *r* denotes samples from rest, and the overline 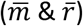 denotes sample mean. *m*∪*r* represents combined movement & rest power sample distributions. *N_m_* and *N_r_* denote the total number of rest and movement samples and *N_m_*∪*_r_* = *N_m_* + *N_r_*. Thus, r^2^ is the percentage of the variance in *m*∪*r* that can be explained by a difference between the individual means in the sub-distributions, 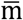 and 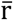. The sign indicates whether power is increasing or decreasing with movement, as illustrated in Fig S4. When viewing figures, consider that all channels were plotted at the interpolated position between the pairs measured, to reflect the change in the brain activity during movement vs. rest. We chose a significance cutoff of one percent of the maximum r^2^ for all channels during a single modality, and insignificant channels were plotted with a white circle of fixed diameter.

#### Timecourse analysis (Fig S7)

sEEG broadband power time series was calculated by 1) band-passing the channel voltage with a 3^rd^ order Butterworth filter in 10Hz bands between 65-115Hz, 2) applying the Hilbert transform and squaring each 10Hz timeseries, and 3) adding the 10Hz timeseries together. The resulting signal was logged, z-scored, smoothed, exponentiated, and centered at zero (i.e. subtracting 1). EMG signal time-series were band passed from 25 to 400Hz^34,35^ using a 3^rd^ order Butterworth filter, notch filtered (60, 120, 180Hz), enveloped, and rectified. These were then logged, z-scored, smoothed, and exponentiated as in prior work.

### Movement type overlap and selectivity in each channel

#### Selective Activity

Selectivity for a specific movement was calculated for each channel in the following manner. The individual r^2^ values for each movement type were multiplied by *e*^iπ/6^ (hand), *e*^*i*π5/6^ (foot), *e*^*i*π3/2^ (foot) and added together. The amplitude of the resulting complex number defines the selectivity, and the phase angle of the complex number points to the movement (or pair of movements) that the channel is selective for. This is illustrated in the 4^th^ row of figure 2.

#### Shared Activity

We calculated overlap of movement representation for each channel as the geometric mean of the individual r^2^ values of all three movement types: 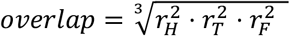. Note that the rare negative r^2^ values (decrease with movement) were set to zero prior to calculation of both selectivity and overlap.

#### Estimation of timing between brain activity and movement (Fig 4)

To estimate the relative latencies between brain activity and movement, we calculated cross correlations between channels of sEEG and EMG signals. These correlations were calculated by taking the dot product of a sEEG channel’s broadband timecourse and the EMG timecourse measuring hand, tongue, and foot movement. Correlations were measured after introducing time delays ranging from −2s to 2s, in 1 sample (.83ms) intervals, obtaining a profile of correlation as a function of latency between the two signals (i.e. a “sliding window” to calculate correlation).

## Code and data availability

All data recorded and code to perform analyses & reproduce the illustrations are publicly available at: https://osf.io/p5n2k

## SUPPLEMENTAL MATERIAL

**Table S1.** Subject Information: Age, Sex, Number of Leads/Laterality, Data Quality, EMG modalities with useful signal, coverage of selective tissue in M1, seizure foci location. *Noise contaminated EMG signal in initial trials; ^◇^Patient refused to have chin EMG stickers placed; ^†^S12 and S13 had low quality data; ^♪^R language dominant

| Subject ID | Age | Sex | # Leads | EMG Recorded | Selective Coverage | <u>Seizure Foci and Lesional Notes</u> | Handed |
| --- | --- | --- | --- | --- | --- | --- | --- |
| S1 | 17 | F | 13/R | H,T,F | H | **Lesion:** R temporal low grade glioma<br>**SOZ:** Right mesial temporal (amygdala, hippocampal body and post. hippocampus), right middle temporal gyrus and right insula | R |
| S2 | 16 | F | 14/L | H,T,F* | H T F | **Lesion:** No lesions<br>**SOZ:** Operculum insula and post central gyrus | R |
| S3 | 16 | M | 15/B | H,T,F | H,F | **Lesion:** Focal Cortical Dysplasia (FCD) in frontal lobe<br>**SOZ:** Frontal lobe near FCD | L |
| S4 | 15 | F | 10/L | H,T,F | H T | **Lesion:** Left mesial temporal lobe sclerosis **SOZ:** L mesial temporal lobe | R |
| S5 | 16 | F | 14/R | H,T,F | H T F | **Lesion:** No lesion<br>**SOZ:** Right precentral postcentral gyri and paracentral lobule | L |
| S6 | 13 | M | 13/R | H,F<sup>◇</sup> | H F | **Lesion:** No lesions<br>**SOZ:** Right temporal | R |
| S7 | 20 | M | 12/R | H,T,F | H T | **Lesion:** Right posterior temporal lobe (prior resection)<br>**SOZ:**<br>1 Posterior -Par-Occ-Temp junction (previous resection)<br>2 Anterolateral Temporal Lobe<br>3 Mesial Temporal Lobe | R |
| S8 | 14 | F | 15/L | H,T,F | N/A | **Lesion:** Left inferior frontal, cortical thickening and blurring of grey-white junction at base of gyrus rectus/orbital gyrus<br>**SOZ:** Left anterior insula/ inferior frontal junction | R |
| S9 | 18 | M | 14/R | H,T,F | H F | **Lesion:** Right mesial temporal lobe sclerosis<br>Anterior temporal lobectomy<br>**SOZ:** Right posterior temporal lobe and insula | L<sup>♯</sup> |
| S10 | 18 | F | 15/L | H,T,F | N/A | **Lesion:** Left occipital and posteromedial temporal: progressed atrophy, cortical/subcortical calcification, and leptomeningeal enhancement.<br>**SOZ (15 seizures):** Left temporo-parieto-occipital region (9), diffuse (5), posterior temporal (1) | R |
| S11 | 11 | M | 12/L | H,T,F | H T | **Lesion:** No lesions<br>**SOZ:** Paracentral Lobule | R |
| S12<sup>+</sup> | 12 | M | 15/R | H,T,F | H | **Lesion:** Posterior paramedian<br>**SOZ multifocal:** Inf occipital, lingual gyrus, hippocampal | R |
| S13<sup>+</sup> | 13 | M | 11/B | H,T,F | H | **Lesion:** No lesions<br>**SOZ:** Right frontal region seizures are generalized. Lennox Gastaut | R |

**Figure S1.**
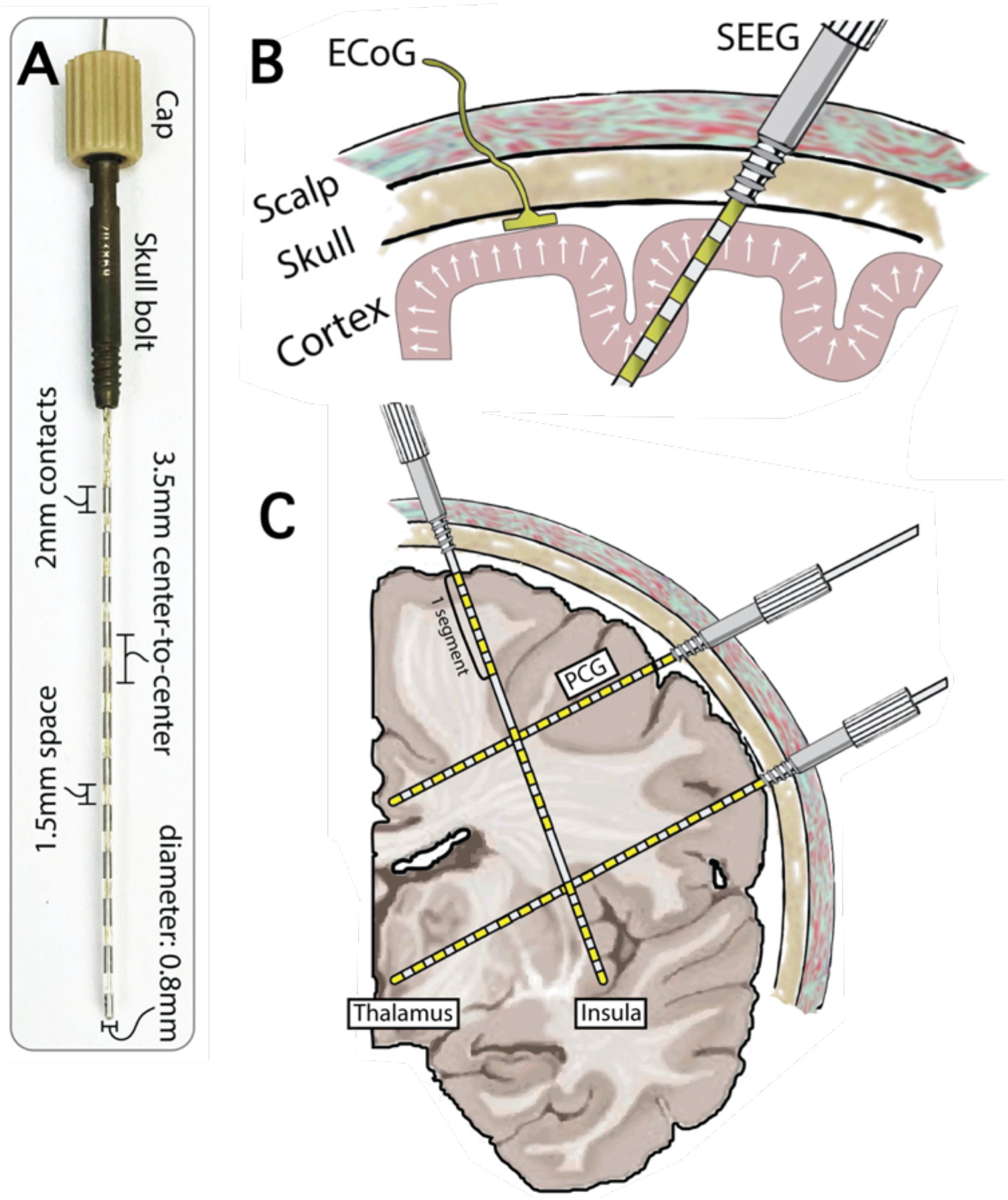
Configuration of sEEG electrodes. **A.** Labeled specifications of sEEG electrodes implanted into subjects. **B.** Schematic illustrating the sulcal coverage provided by sEEG and the limitation of ECoG electrodes to the brain’s contour. **C.** Hemispheric coronal slice showing segmented and non-segmented sEEG leads with the ability to record from the entire brain volume.

**Figure S2.**
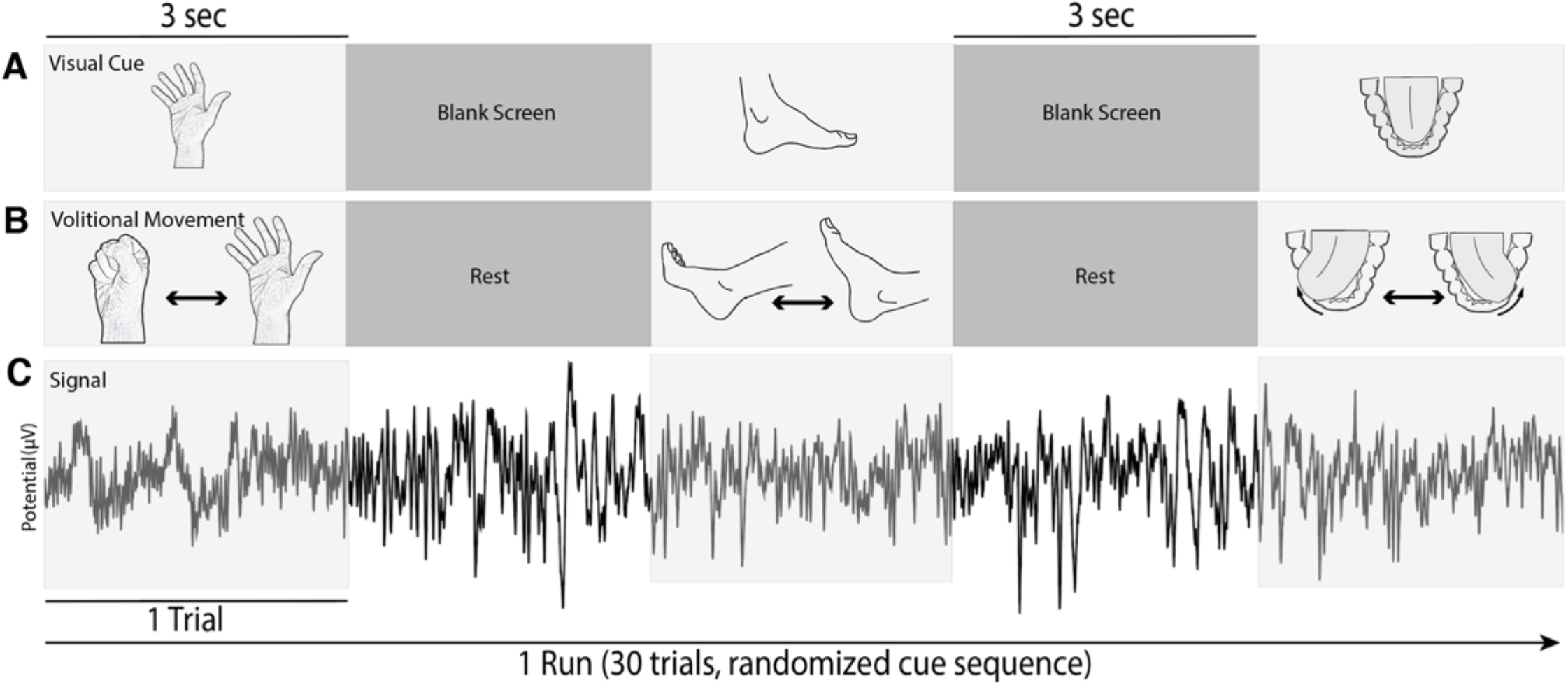
Motor Task Design. **A.** Patients were presented with a 3 second visual cue on a screen 75-100 cm away displaying either a hand, foot, or tongue with a blank screen interleaved. **B.** Subjects were instructed to open and close their hand, dorsi and plantar flex their foot, and move their tongue laterally with mouth closed upon visualization of cue of hand, foot, tongue respectively. Subjects were instructed to remain still when the screen was blank. **C.** sEEG signal was recorded across all trials and the entire set of trials (hand, foot, or tongue movement periods) in a single run. There were 30 trials per run.

**Figure S3.**
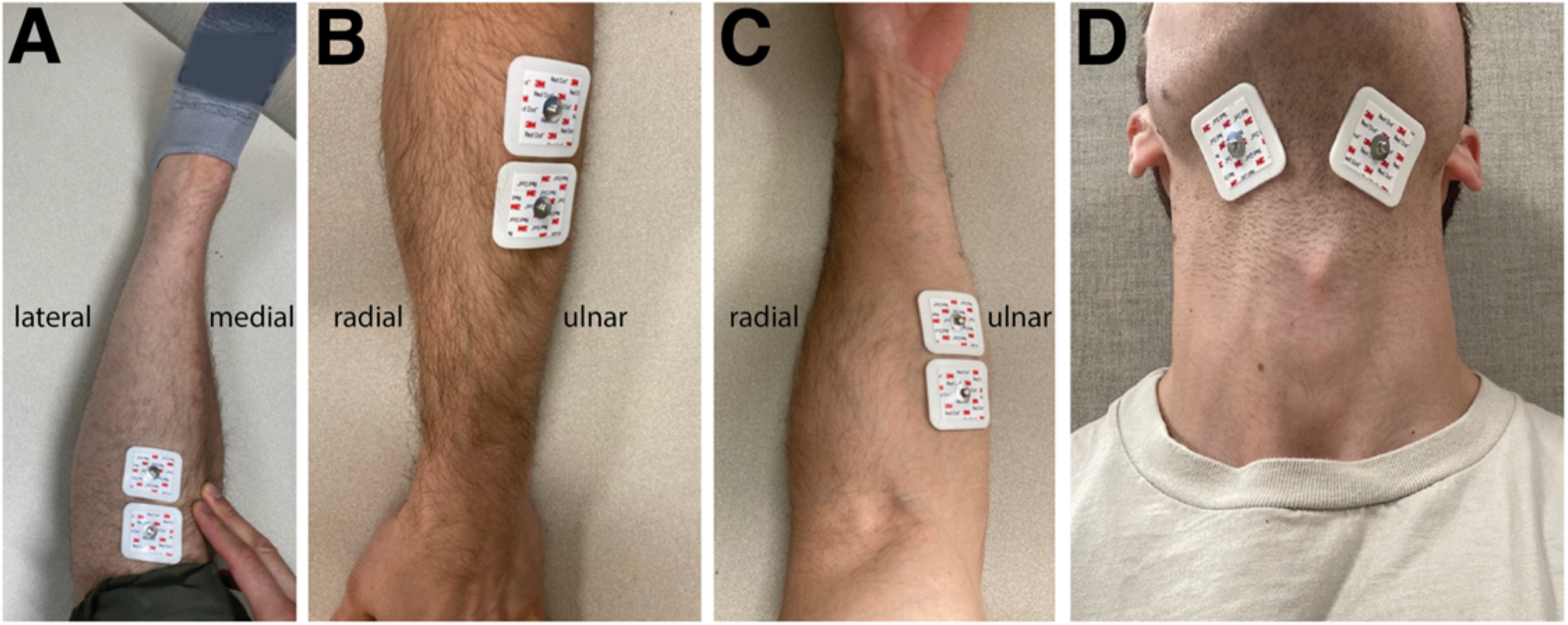
EMG Sticker Placement. **A.** Foot (i.e. anterior tibialis) **B.** Hand (i.e. extensor carpi ulnaris, extensor digitorum) **C.** Forearm Flexors (i.e. flexor carpi radialis). **D.** Tongue (i.e. suprahyoid muscles).

**Figure S4.**
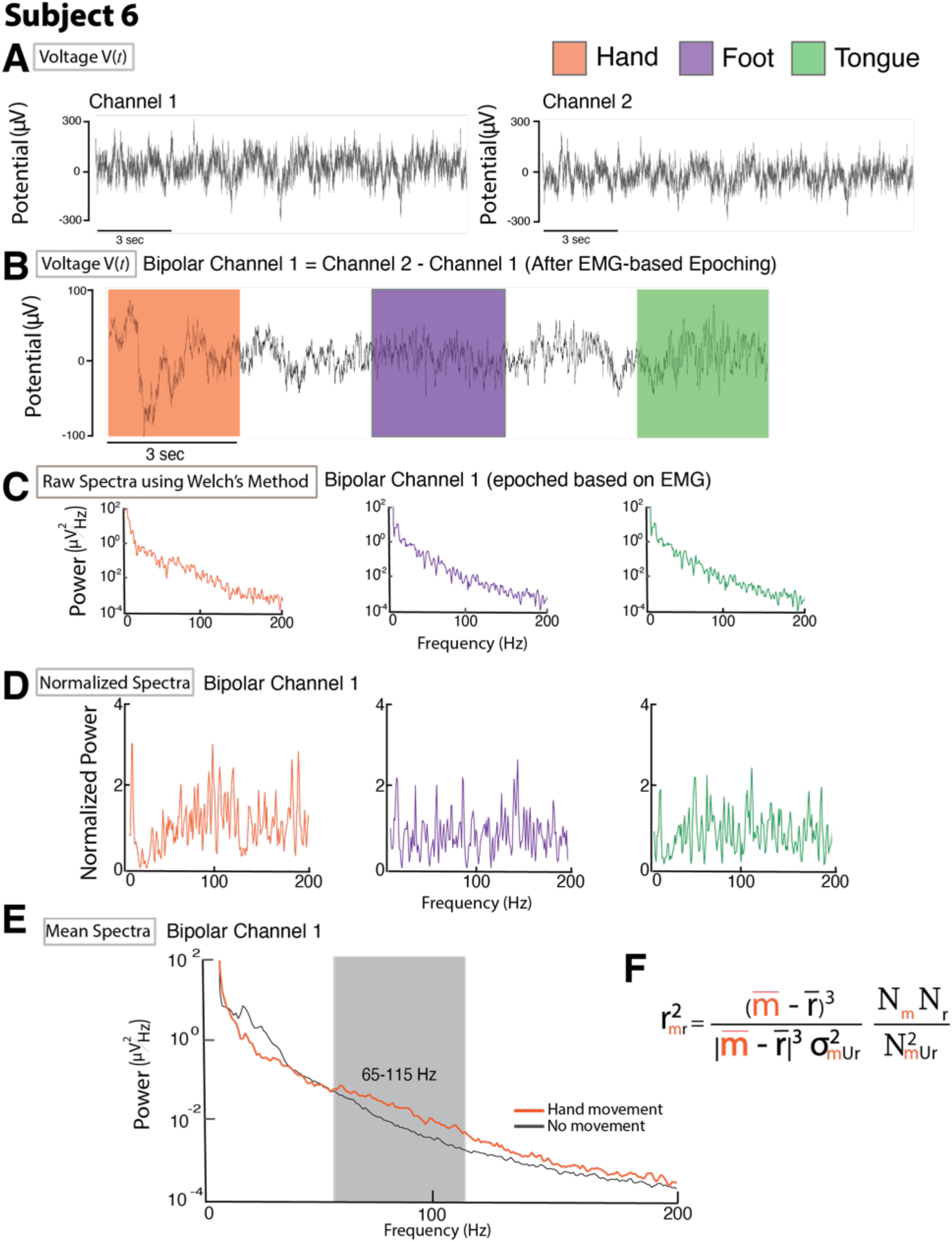
Basic Spectral Calculations. **A.** Adjacent channels within the same segment of the same lead were re-referenced in a bipolar fashion. **B.** Voltage time series were segmented using EMG-defined movement periods. **C.** The power spectral density of each trial was calculated. **D.** Spectra were normalized to the mean spectra and logged. **E.** Average power spectra for all trials of the same movement type (e.g. mean of hand trials) were used to compare 65-115Hz (HFB) power between movement and rest. **F.** r^2^ values were calculated using the mean HFB power during trials of a single movement type and the rest periods immediately following.

**Figure S5.**
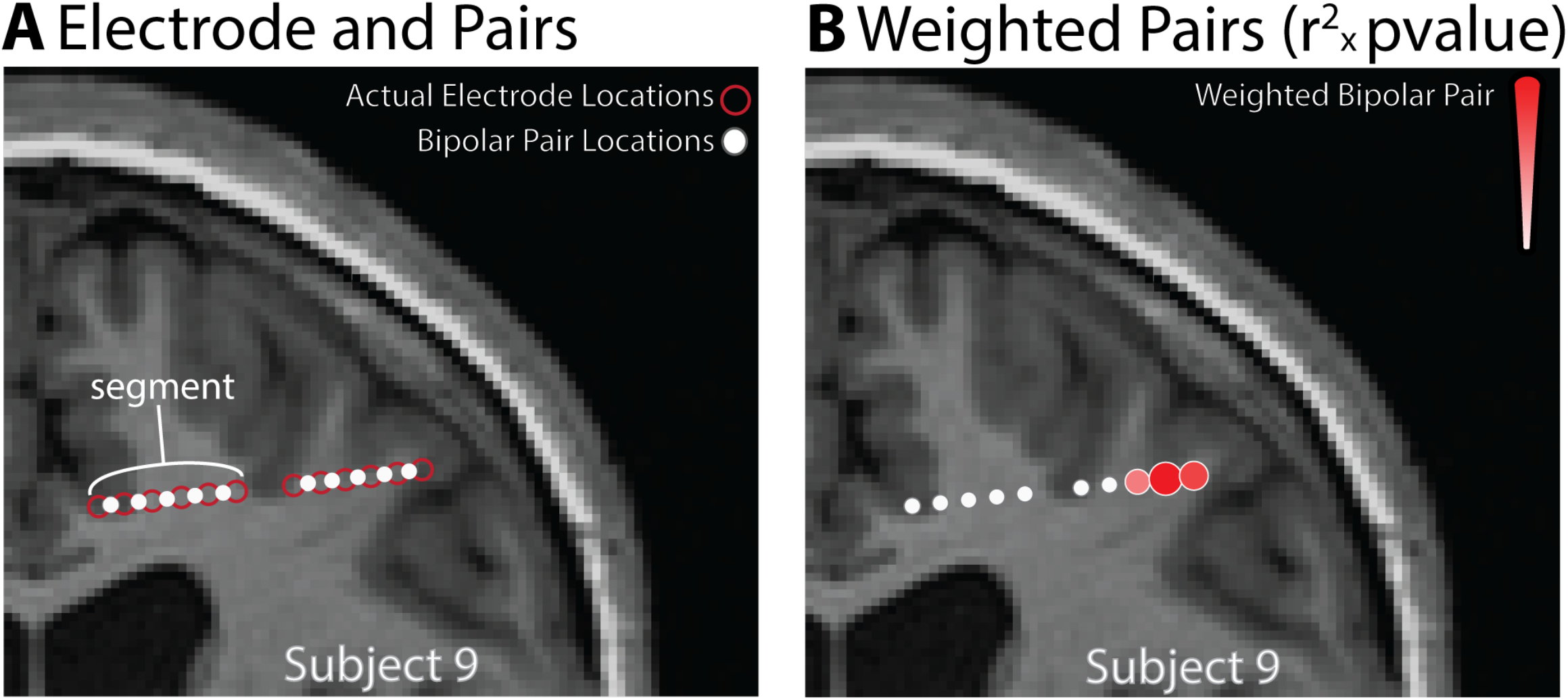
Electrode plots reflect bipolar pairs and r^2^ values. **A.** Plotted circles represent the interpolated points between the two electrode channels making up a differential pair. **B.** The sign and magnitude of the r^2^ correlation coefficient determines the size and color hue of each electrode pair plotted.

**Figure S6.**
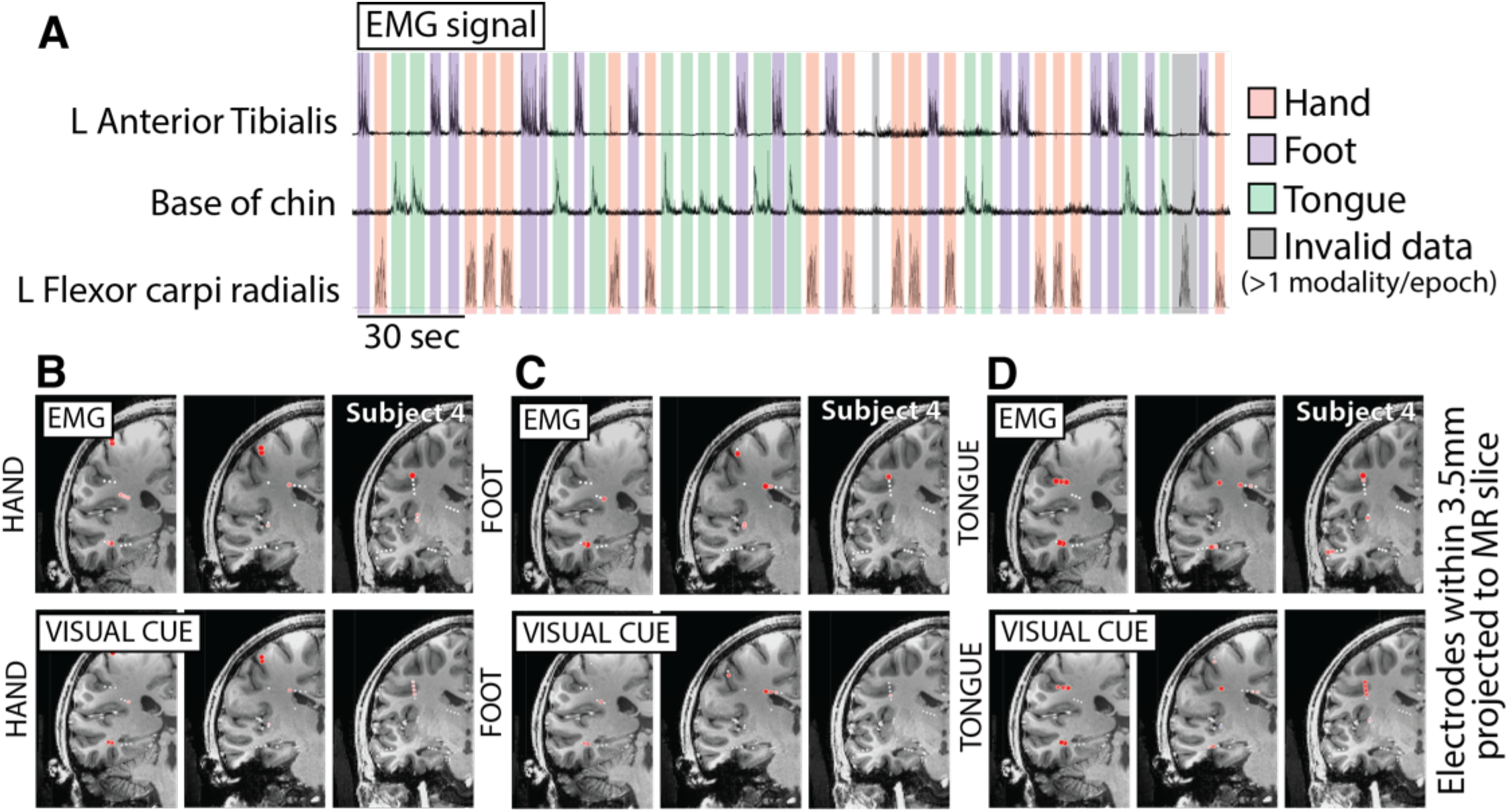
EMG vs. cue-based segmentation. **A.** EMG signals measured between pairs of channels placed as seen in Figure S3. Trials specific to hand, foot, or tongue are shaded in orange, purple, and green respectively with gray demonstrating epochs with simultaneous movement of multiple modalities. The absence of background shading indicates resting. **B-D.** Comparison of results between EMG and stimulus-based data segmentation for hand, foot, and tongue movement in Subject 4. Note that maps differ between the two segmentation methods.

**Figure S7.**
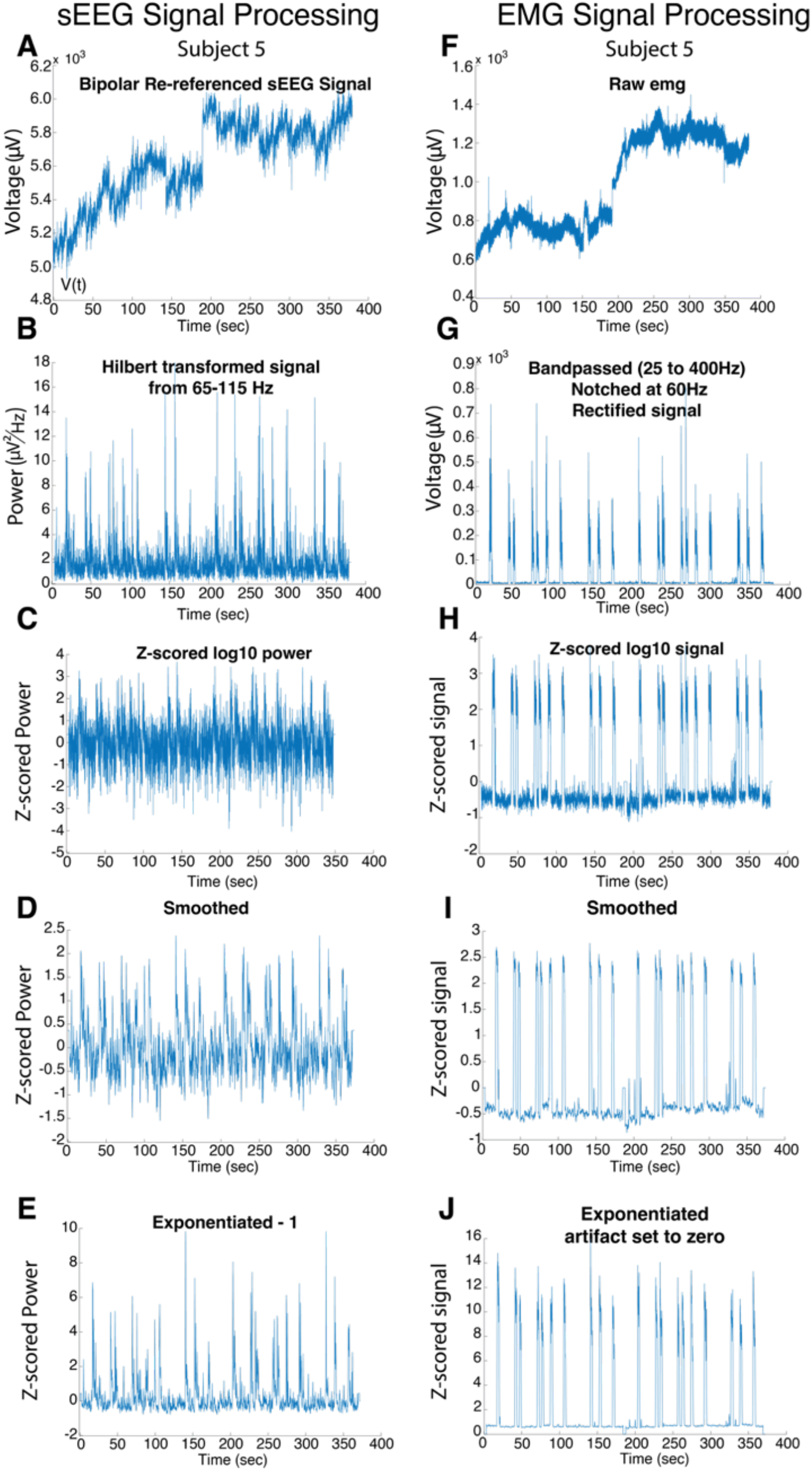
Time Series Signal Processing. **Subject 5 A.** The sEEG signal were bipolar re-reference **B.** The sEEG signal was band-passed using a 3^rd^ order Butterworth filter in 10Hz bands (from 65-115Hz), Hilbert transformed {Hilbert(V(t)) = V(t) + iH(t) = re^iΩ^}, squared, and the 10Hz timeseries were added together. The signal was then **(C)** logged and z scored, **(D)** smoothed with a moving average window of 500ms, and then **(E)** exponentiated and centered around zero by subtracting 1. **F.** The EMG was recorded in a bipolar fashion, then **(G)** band-passed from 25 to 400Hz, **(H)** logged and z scored, **(I)** smoothed with a moving average window of 500ms, and then **(J)** exponentiated. Inter-run artifact was set to zero.

**Figure S8.**
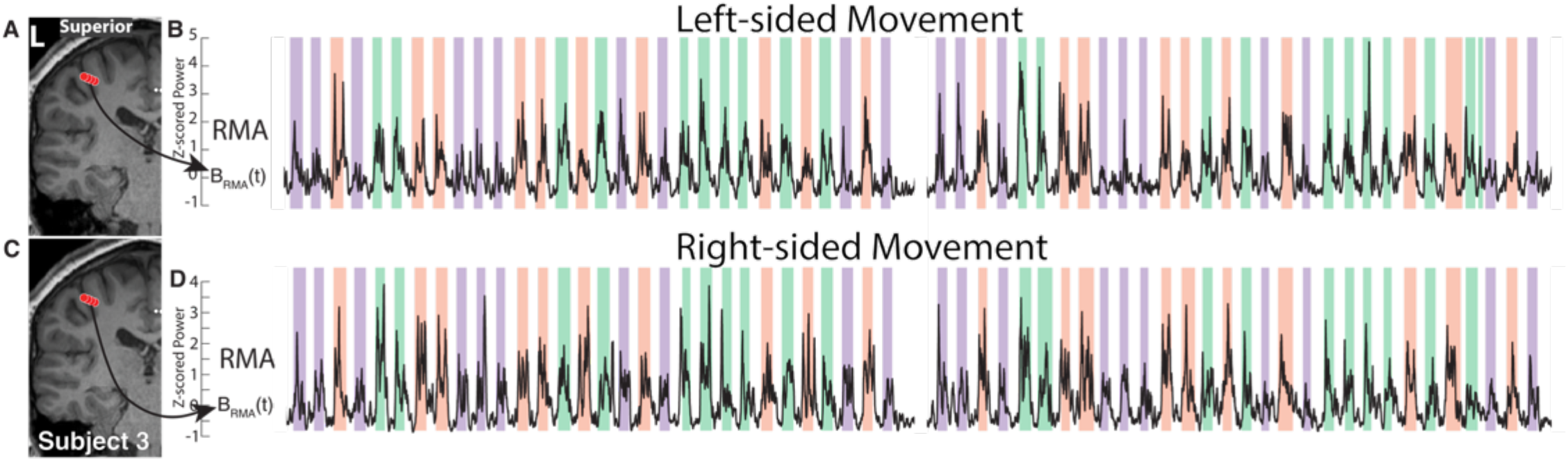
RMA activity for movements of both sides of the body. **A.** Electrodes within RMA during right-sided movement of hand, tongue, foot **B.** Time-series of broadband power across two consecutive runs from the RMA site during left-sided movement **C.** Electrodes within RMA during right-sided movement of hand, tongue, foot **D.** Time-series of broadband power across two consecutive runs from the RMA site during right-sided movement.

**Supplemental Figure S9.**
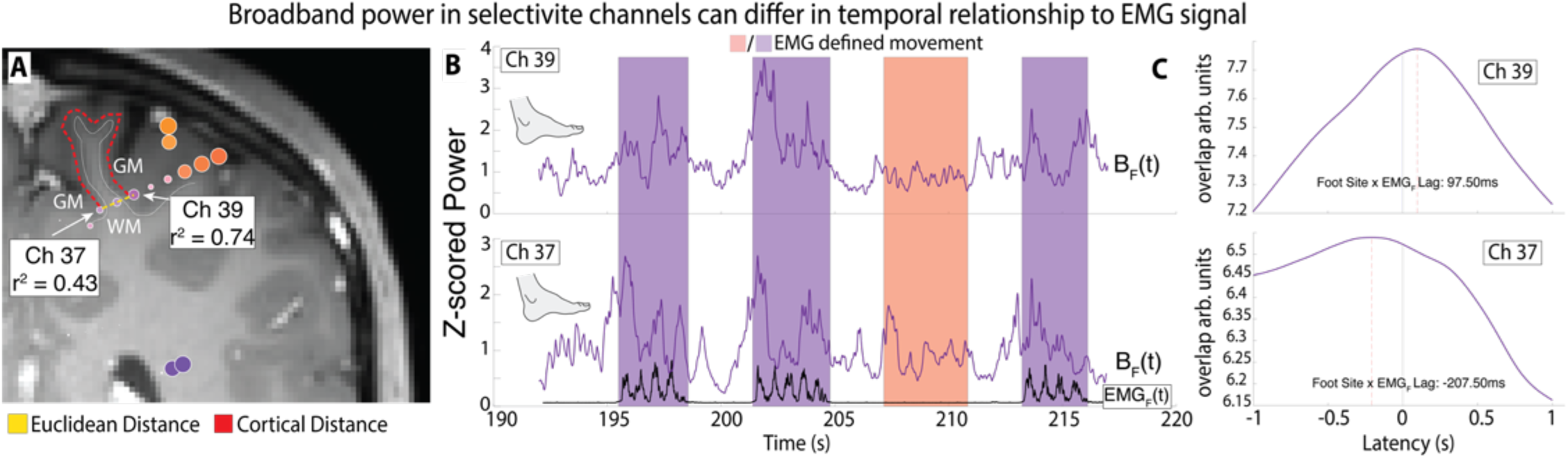
Selective regions differ in relation to EMG onset. **A.** Coronal slice of selective eletrodes for foot (arrows) with small Euclidean distance yet large cortical distance. **B.** Broadband timeseries show a peak at the onset of movement as defined by EMG. **C.** Latency between the two pairs and EMG differ. The pair near the established foot region (Ch 37) peaks prior to EMG foot onset while Ch39 peaks after onset. This might be attributed to the possibility that it is sensory tissue in the posterior aspect of the base of the central sulcus.

**Figure S10.**
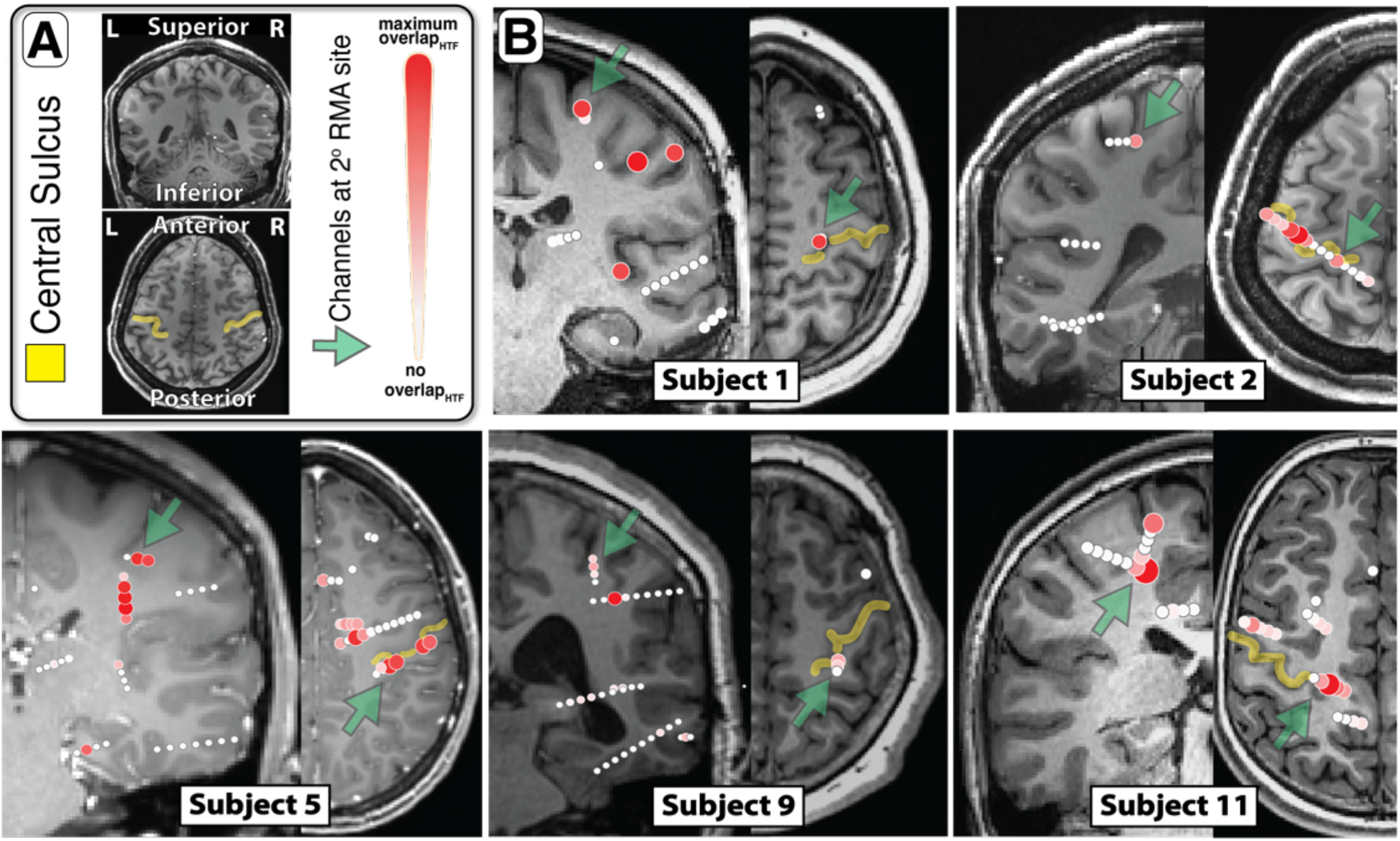
A secondary Rolandic motor association area within the central sulcus. **A.** Orientation of coronal and axial slices **B.** Axial and coronal views demonstrate localization of channels with shared representation between hand, tongue, and foot movement within the central sulcus. Green arrows indicate the channels which lie within the central sulcus at a secondary RMA site, superior and medial to the primary RMA site found in every patient.

**Supplemental Figure S11.**
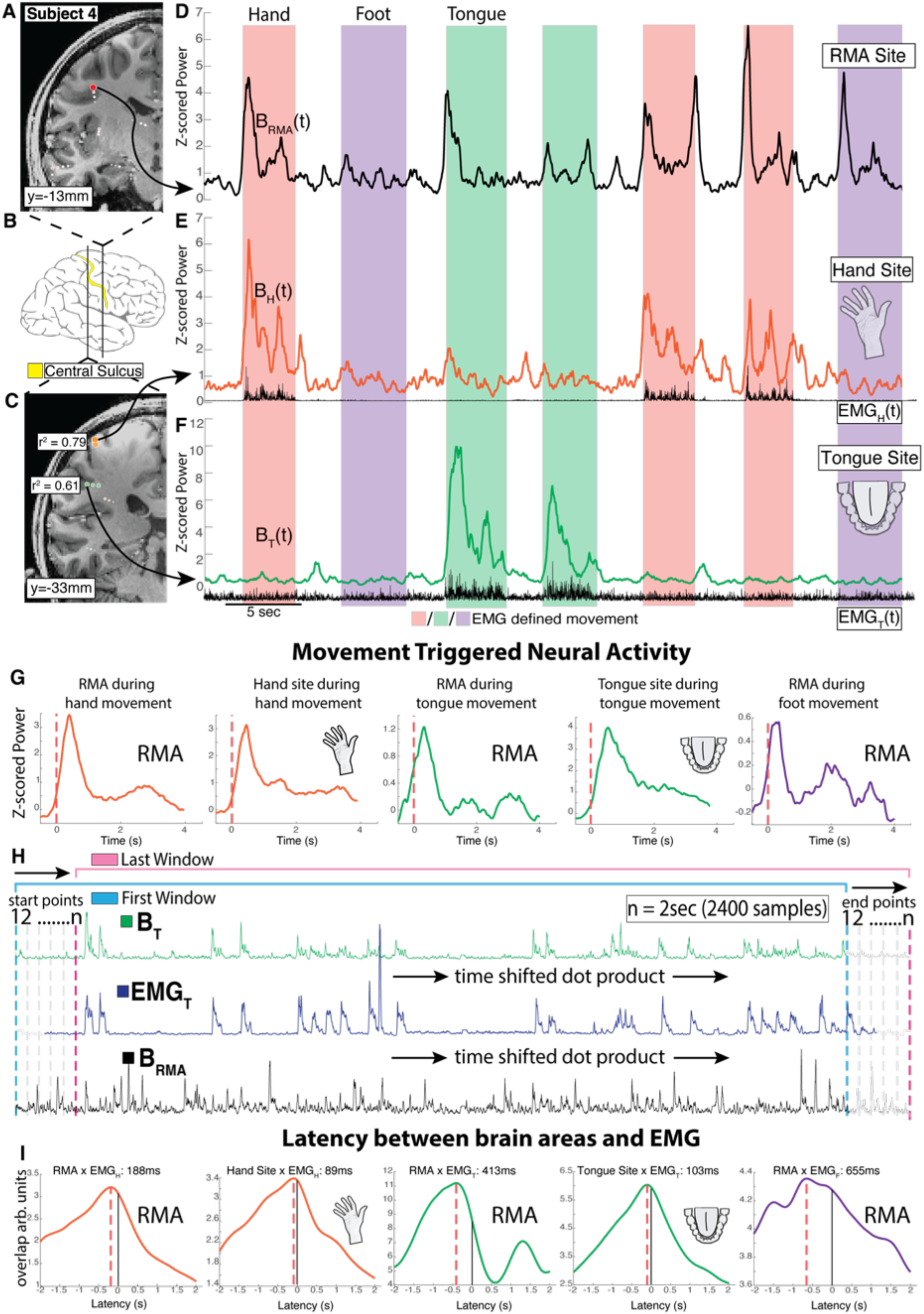
Temporal dynamics of neural activity in the precentral gyrus, subject 4. **A.** Coronal T1 MRI cross section through the precentral gyrus (PCG) and central sulcus in the left hemisphere, with plotted shared activity revealing the RMA area, as in Fig 2H. **B.** Position of coronal slices in (A)&(C). **C.** Left hemisphere, with plotted selectivity revealing somatotopic hand and tongue representation, as in Fig 2I. **D-F.** Timecourse of broadband activity (65-115Hz, “B”) for RMA, foot, hand channels, reflecting local neural activity. Background shading indicates EMG-defined movement periods of the hand (red), foot (purple), and tongue (green). **G.** Brain activity averaged to onset of foot, hand, and tongue EMG. **H.** Schematic of time-shifted dot products to calculate latencies between brain activity and movement. **I.** Profiles of latency between brain areas (RMA, tongue selective, hand selective) and EMG (tongue and hand).

**Supplemental Figure S12.**
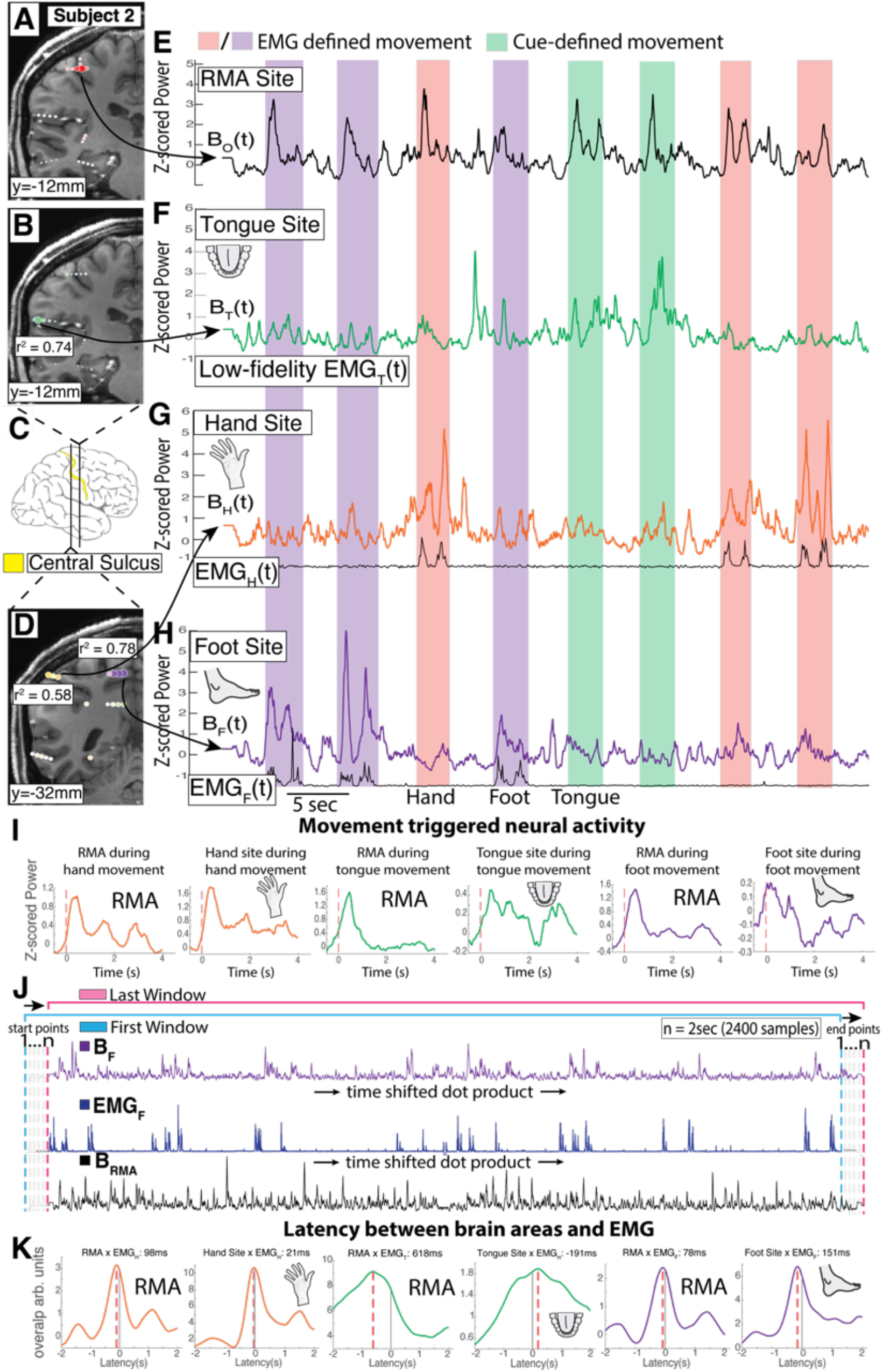
Temporal dynamics of neural activity in the precentral gyrus, subject 2. **A.** Coronal T1 MRI cross section through PCG in the left hemisphere, with plotted shared activity revealing the RMA area, as in Fig 2H. **B.** Left hemisphere, with plotted selectivity revealing somatotopic tongue representation, as in Fig 2I. **C.** Position of coronal slices in (A)&(C). **D.** As in (B), with plotted selectivity revealing somatotopic foot and hand representation. **E-H.** Timecourse of broadband activity (65-115Hz, “B”) for RMA, tongue, hand, foot channels, reflecting local neural activity. Background shading indicates EMG-defined movement periods of the hand (red), foot (purple), and tongue (green). **I.** Brain activity averaged to onset of foot, hand, and tongue EMG. **J.** Schematic of time-shifted dot products to calculate latencies between brain activity and movement. **K.** Profiles of latency between brain areas (RMA, hand selective, tongue selective, and foot selective) and EMG (hand tongue and foot).

**Supplemental Figure S13.**
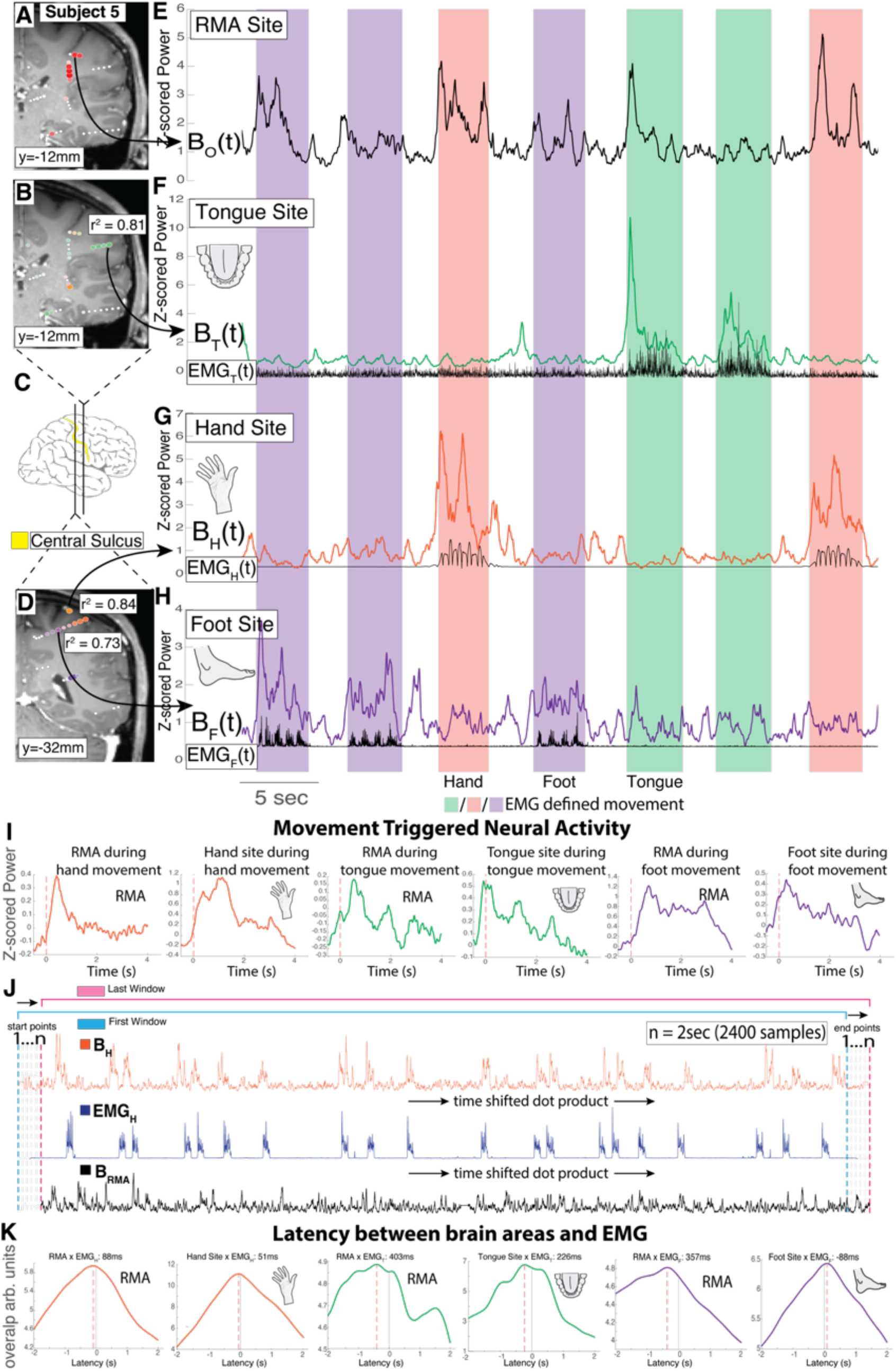
Temporal dynamics of neural activity in the precentral gyrus, subject 5. **A.** Coronal T1 MRI cross section through the precentral gyrus and central sulcus in the right hemisphere, with plotted shared activity revealing the RMA area, as in Fig 2H. **B.** Right hemisphere, with plotted selectivity revealing somatotopic tongue representation, as in Fig 2I. **C.** Position of coronal slices in (A)&(C). **D.** Right hemisphere, with plotted selectivity revealing somatotopic foot and hand representation, as in Fig 2I. **E-H.** Timecourse of broadband activity (65-115Hz, “B”) for RMA, tongue, hand, foot channels, reflecting local neural activity. Background shading indicates EMG-defined movement periods of the hand (red), foot (purple), and tongue (green). **I.** Brain activity averaged to onset of foot, hand, and tongue EMG. **J.** Schematic of time-shifted dot products to calculate latencies between brain activity and movement. **K.** Profiles of latency between brain areas (RMA, hand selective, tongue selective, and foot selective) and EMG (hand tongue and foot).

**Figure S14.**
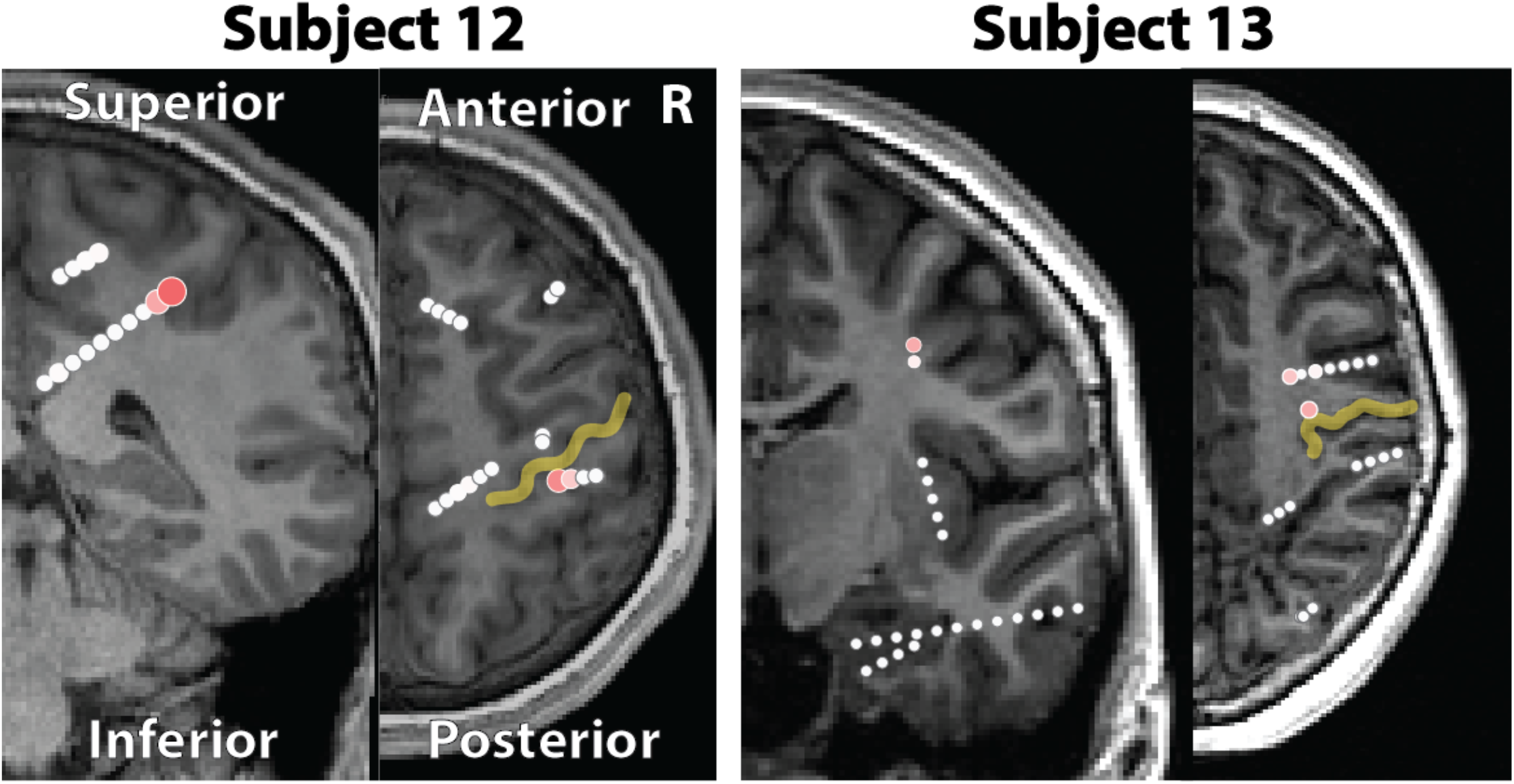
Rolandic motor association area within the central sulcus: subjects with low-fidelity data. Coronal and axial slices of overlapping electrode pairs between hand, tongue, and foot movement for Subjects 10 & 11. Axial and coronal views demonstrate localization of overlapping pairs within the central sulcus. Due to low quality of task compliance (e.g. fidgeting, myoclonic bursts) there were very few quality trials. The results are similar, but the fidelity of these data are low as the trial numbers are few.

**Figure S15.**
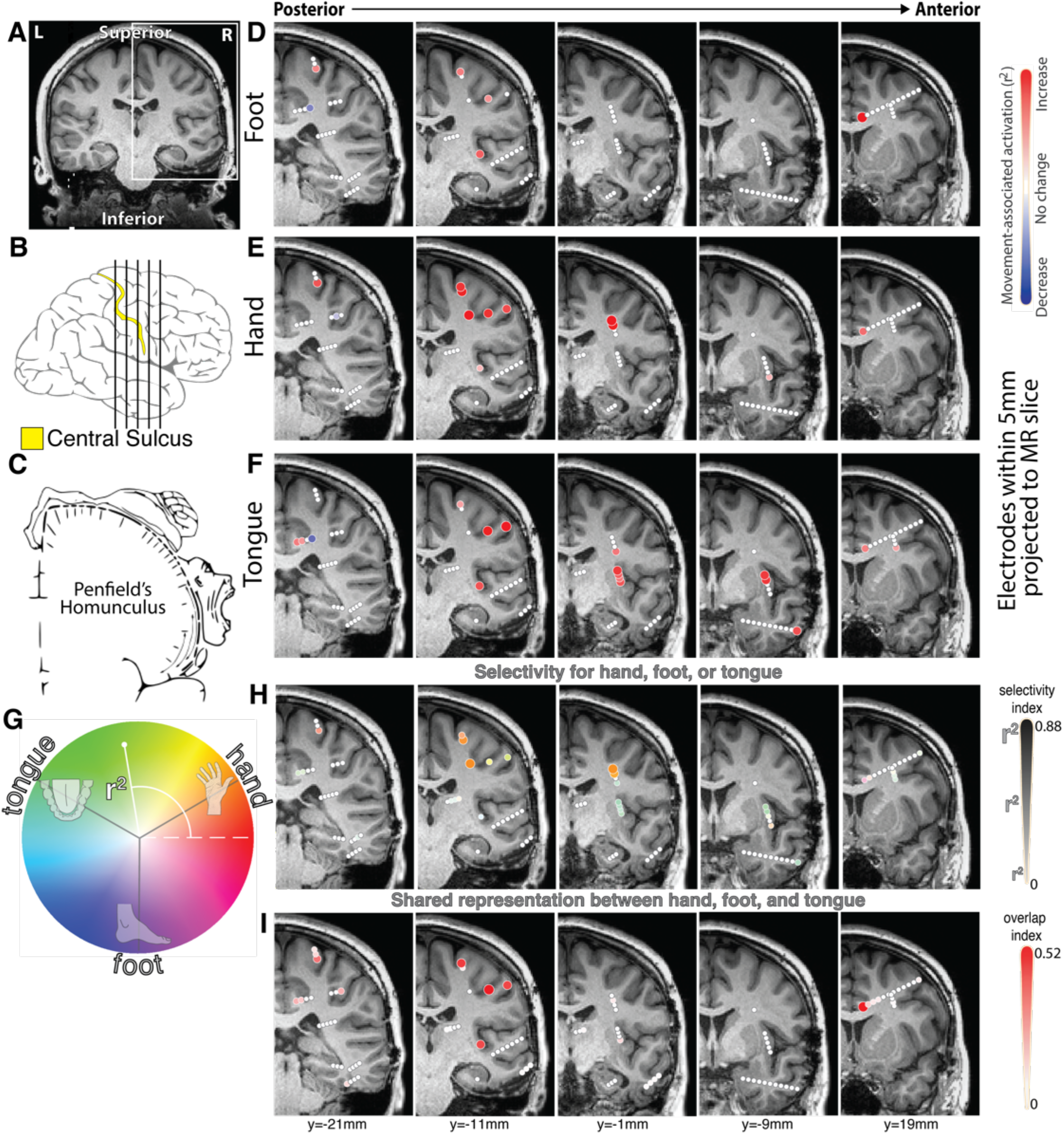
As in the main text Figure 2, for subject 1.

**Figure S16.**
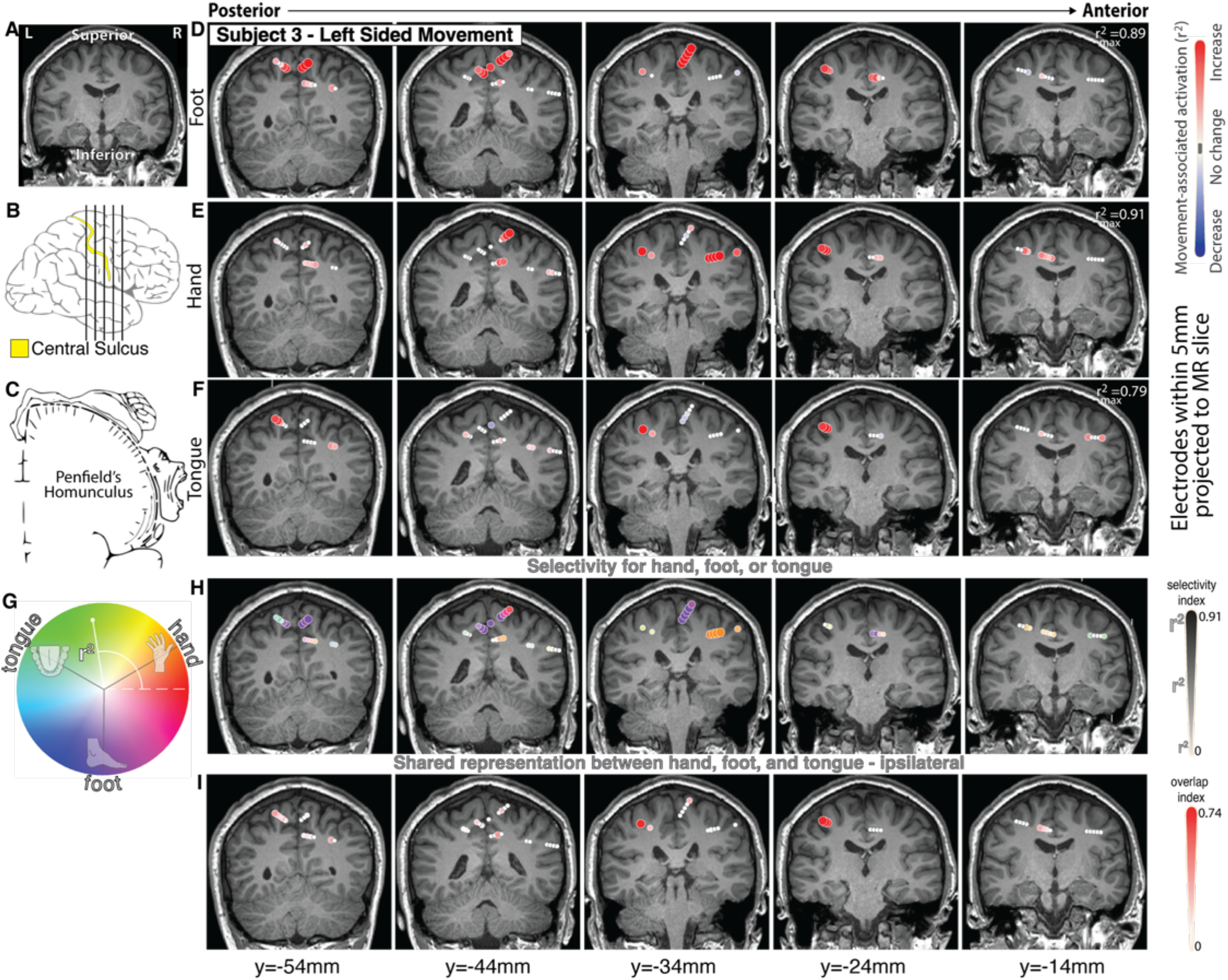
As in the main text Figure 2, for subject 3, during <u>left-sided</u> movement.

**Figure S17.**
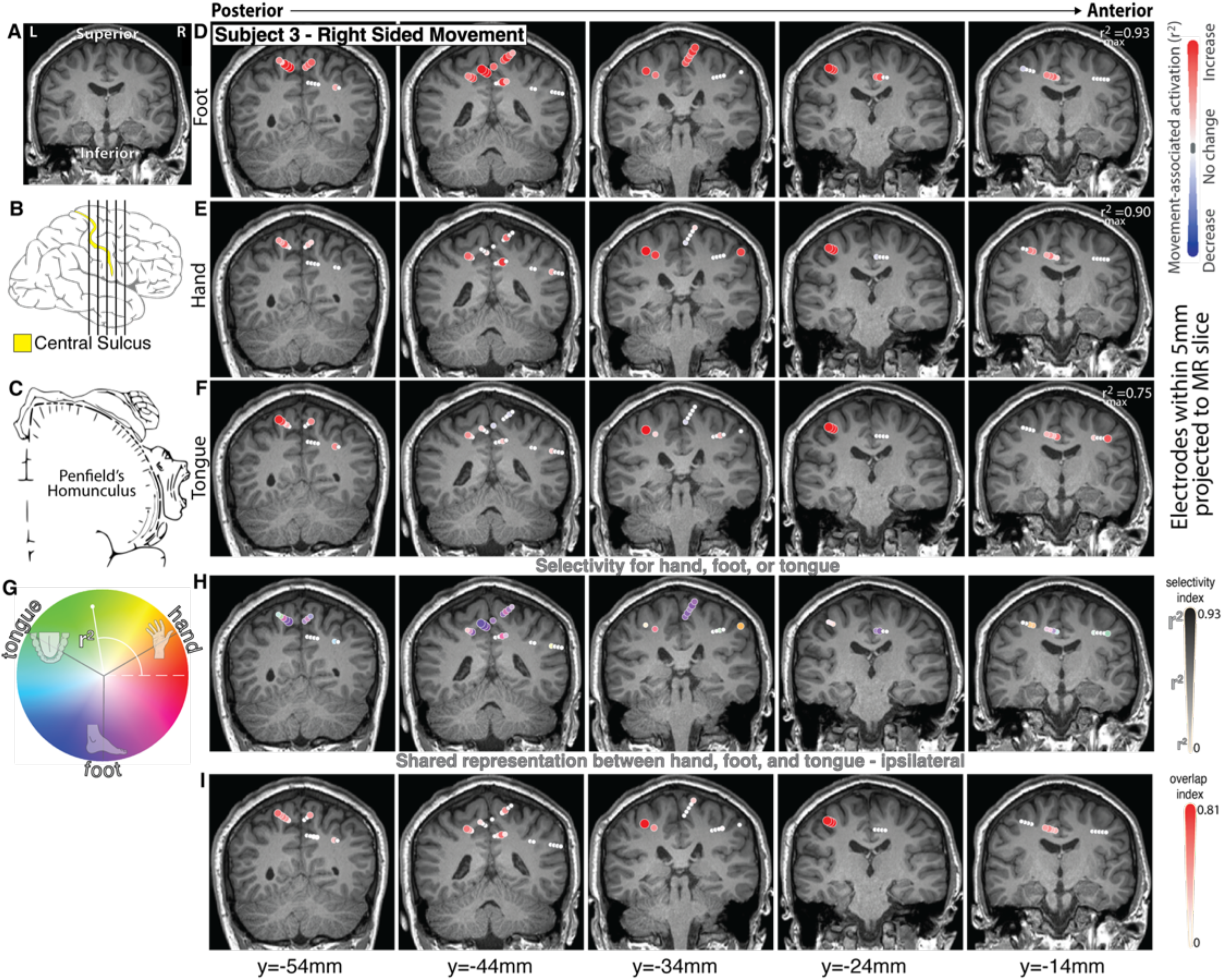
As in the main text Figure 2, for subject 3, during <u>right-sided</u> movement.

**Figure S18.**
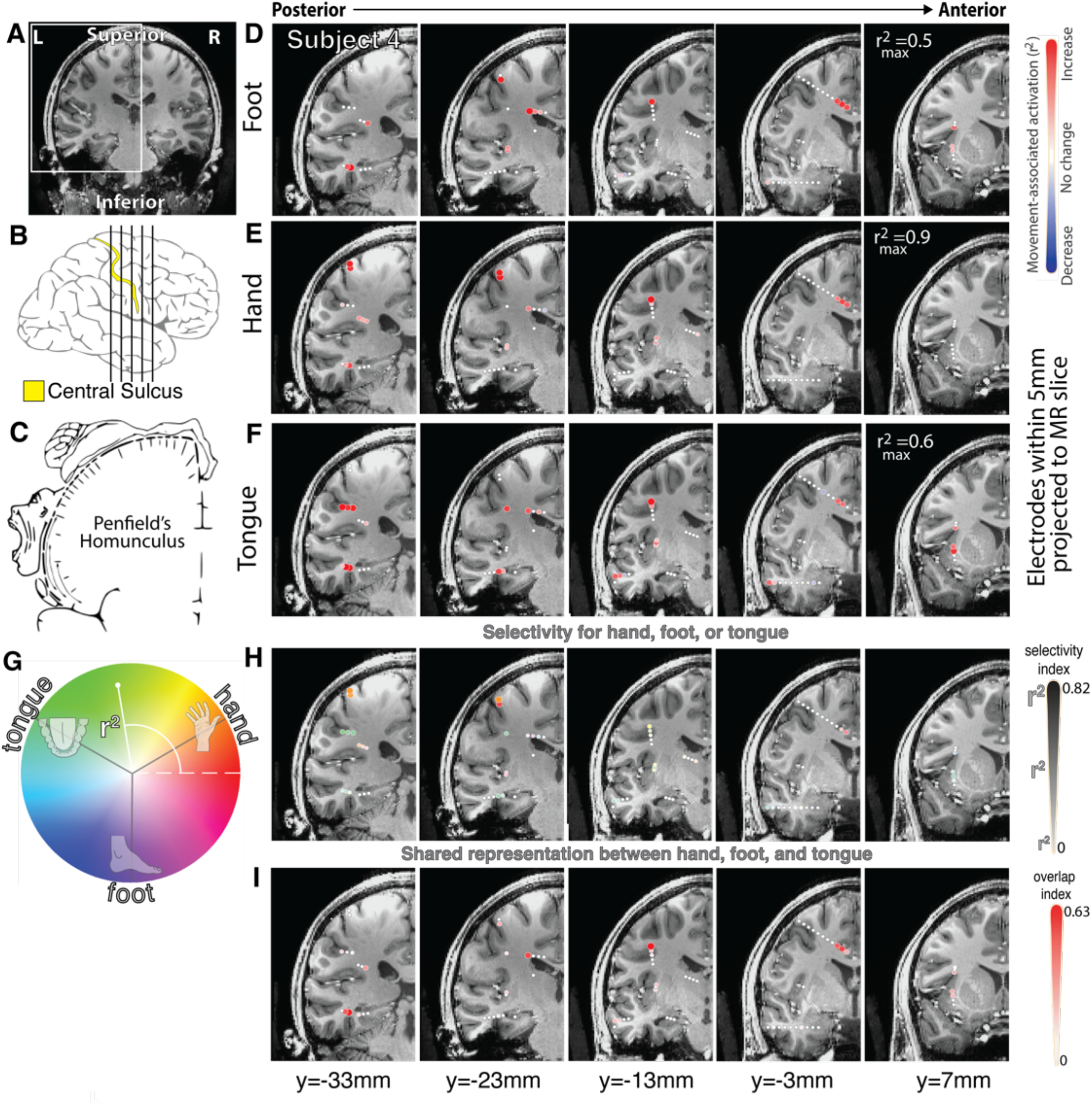
As in the main text Figure 2, for subject 4.

**Figure S19.**
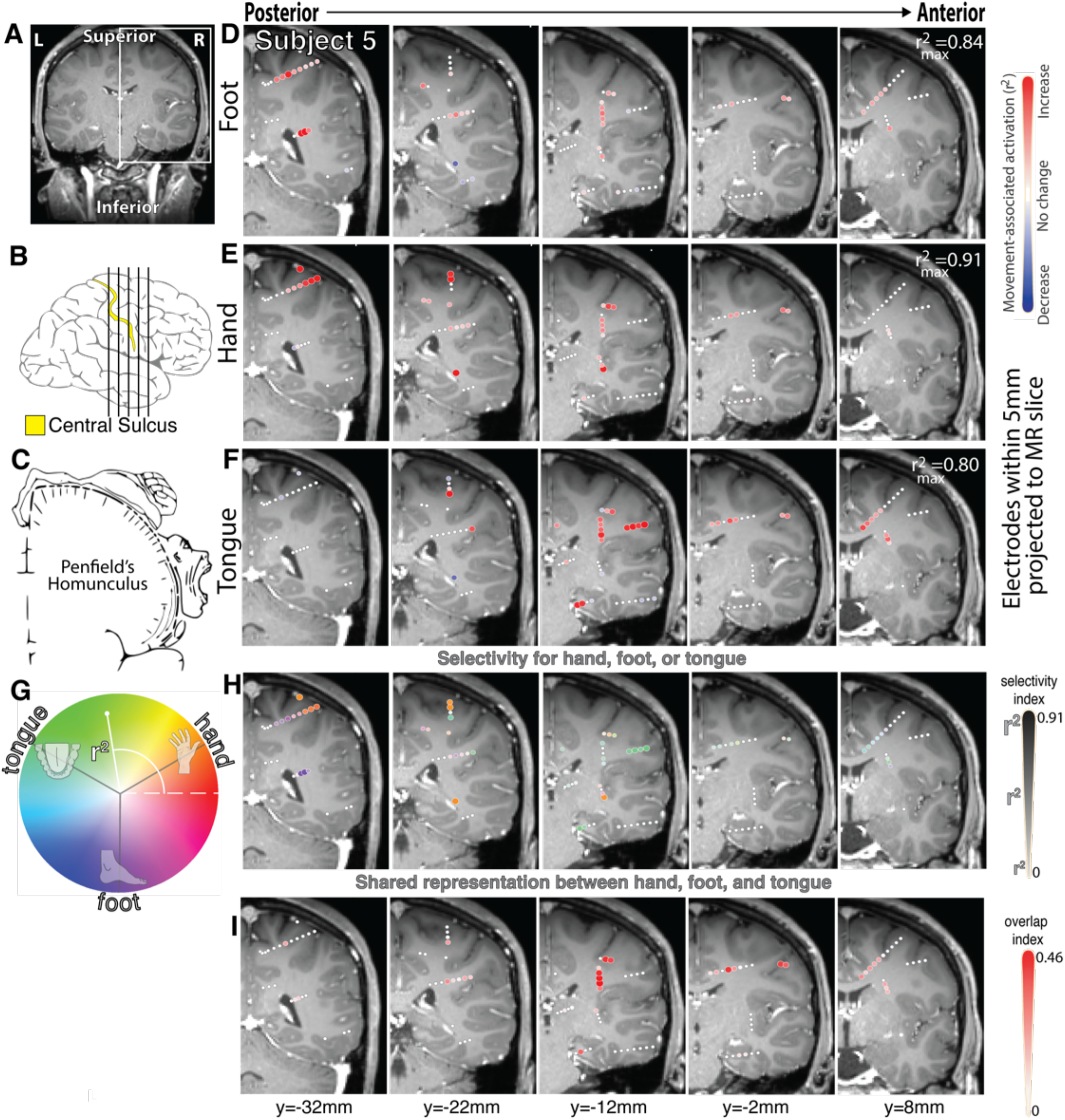
As in the main text Figure 2, for subject 5.

**Figure S20.**
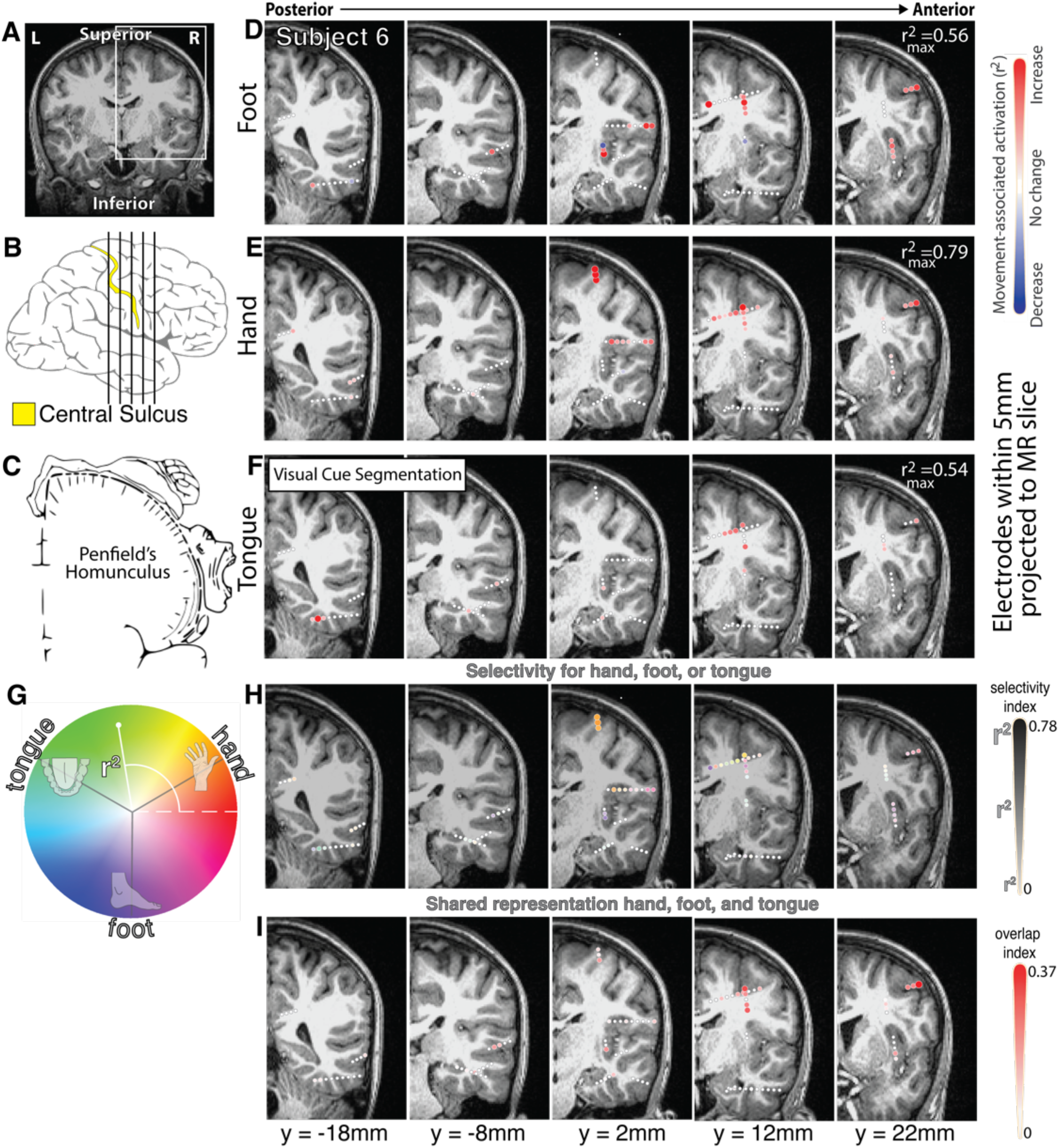
As in the main text Figure 2, for subject 6.

**Figure S21.**
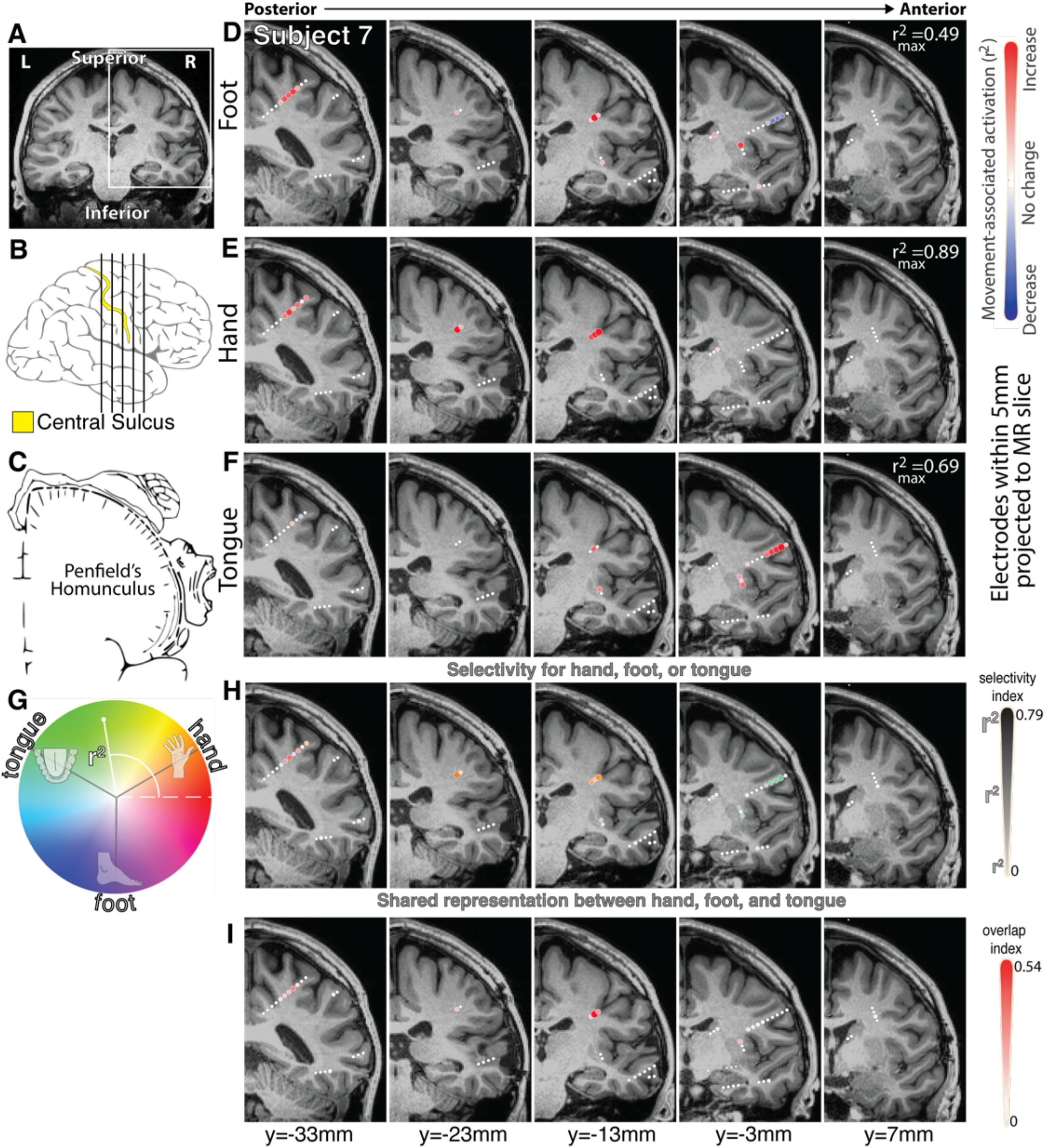
As in the main text Figure 2, for subject 7.

**Figure S22.**
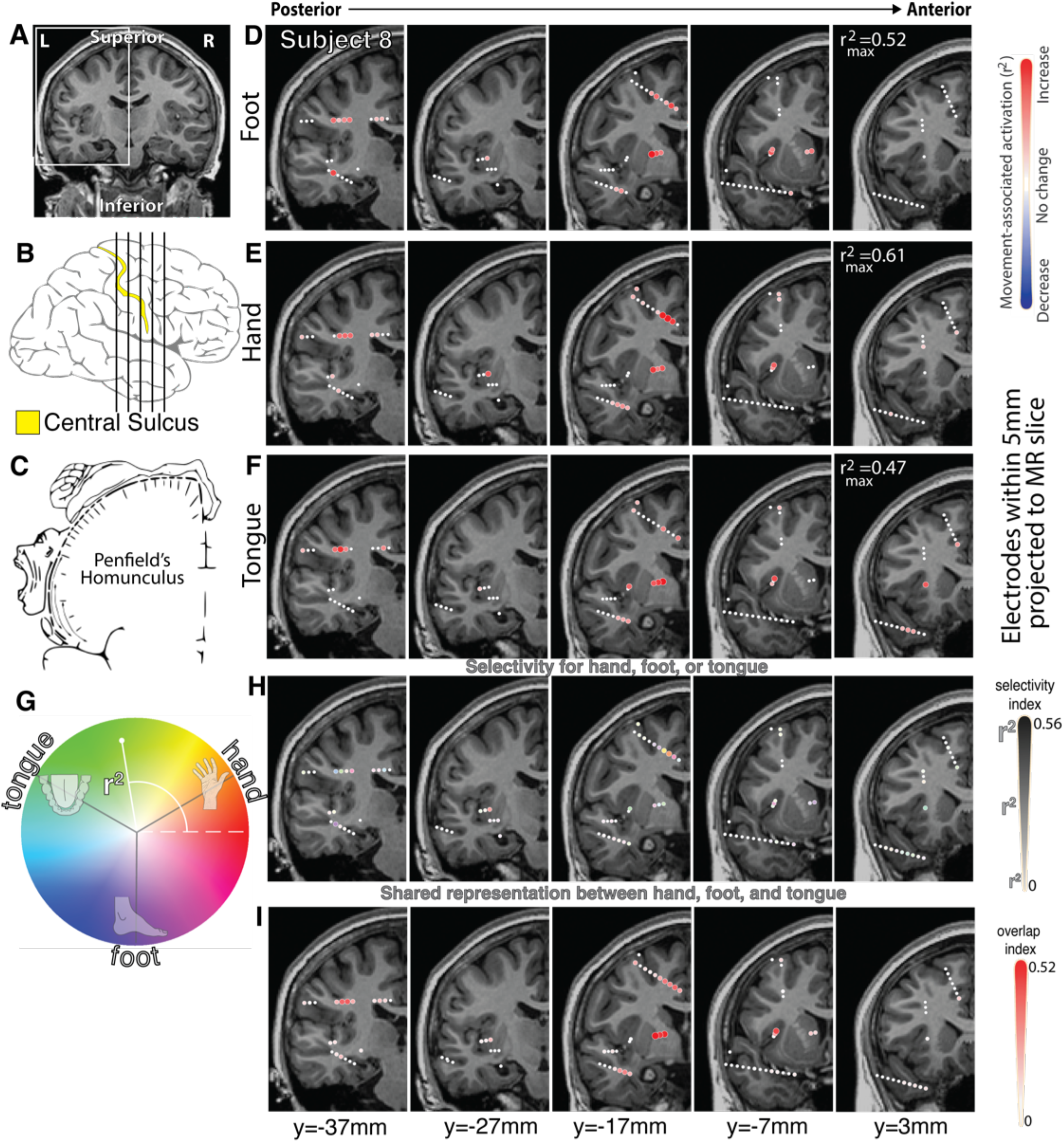
As in the main text Figure 2, for subject 8.

**Figure S23.**
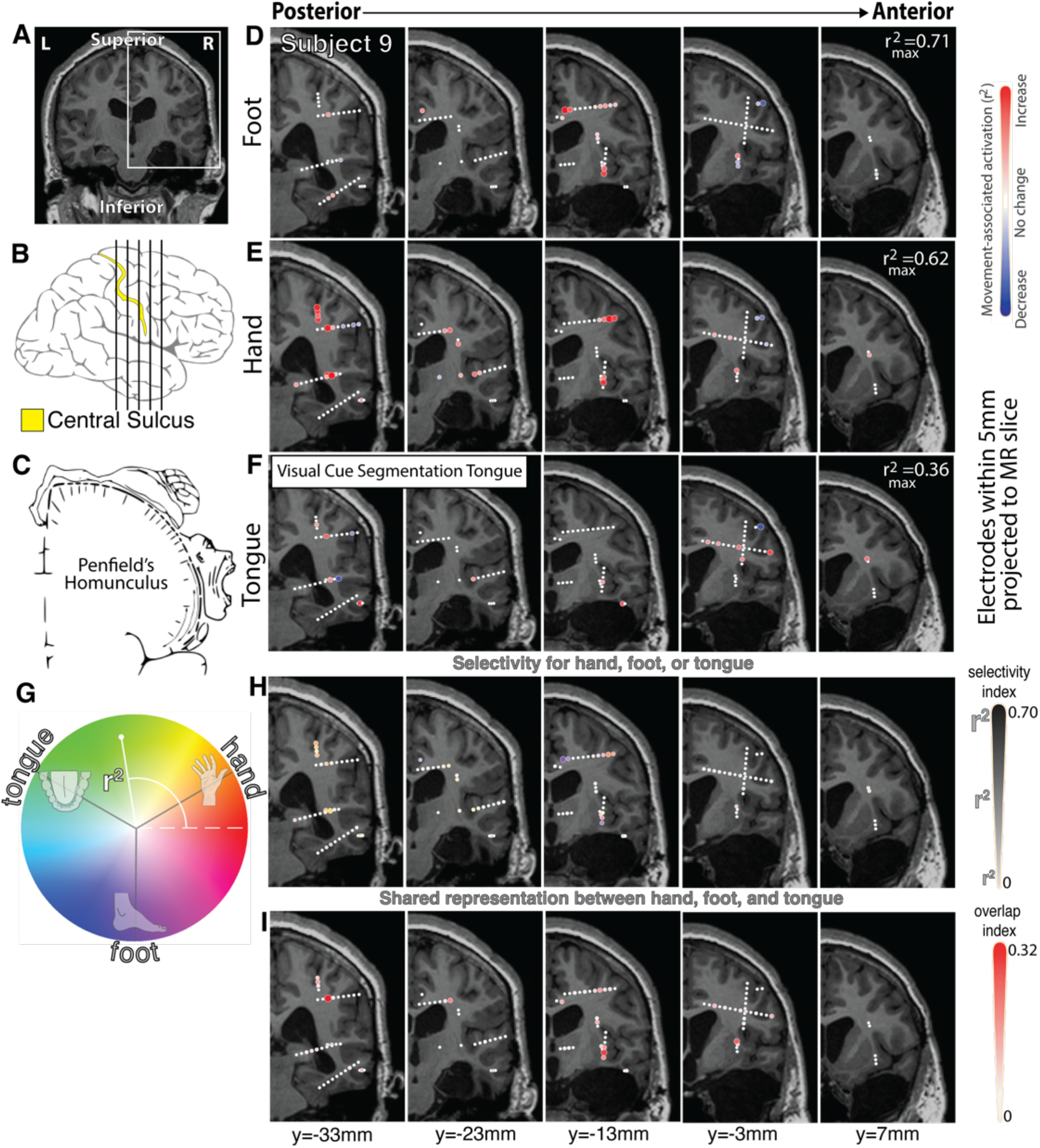
As in the main text Figure 2, for subject 9.

**Figure S24.**
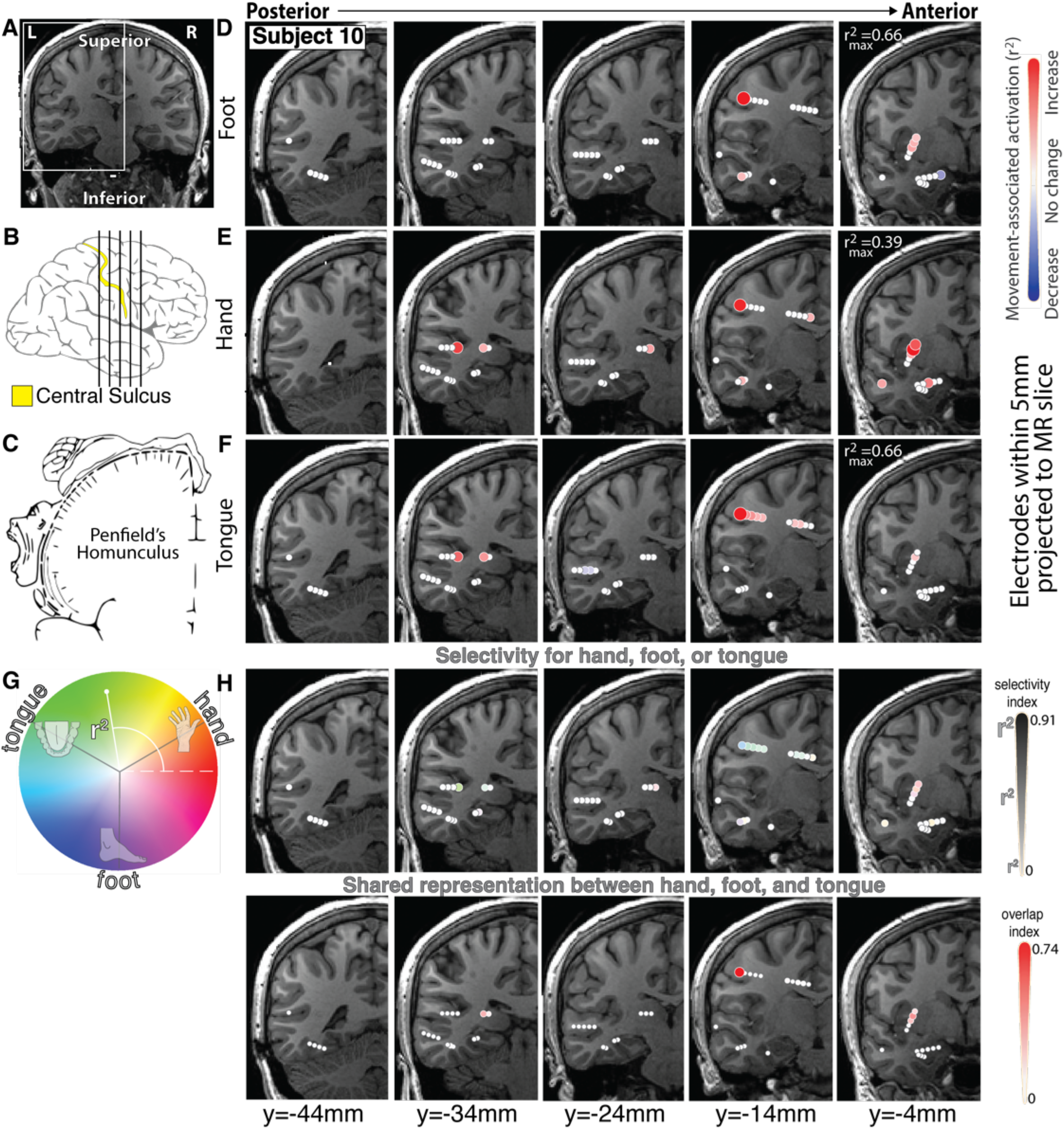
As in the main text Figure 2, for subject 10.

**Figure S25.**
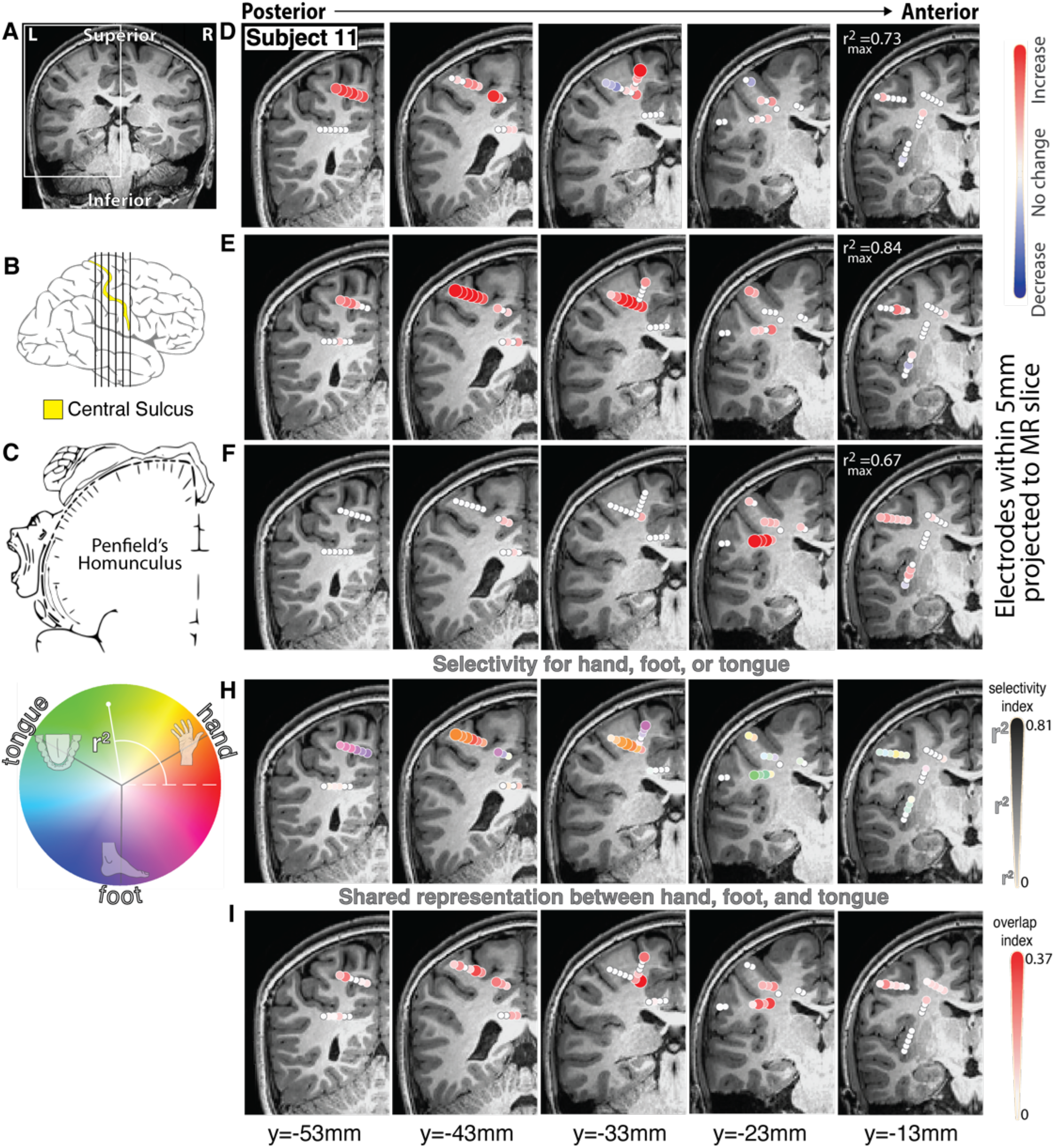
As in the main text Figure 2, for subject 11.

**Figure S26.**
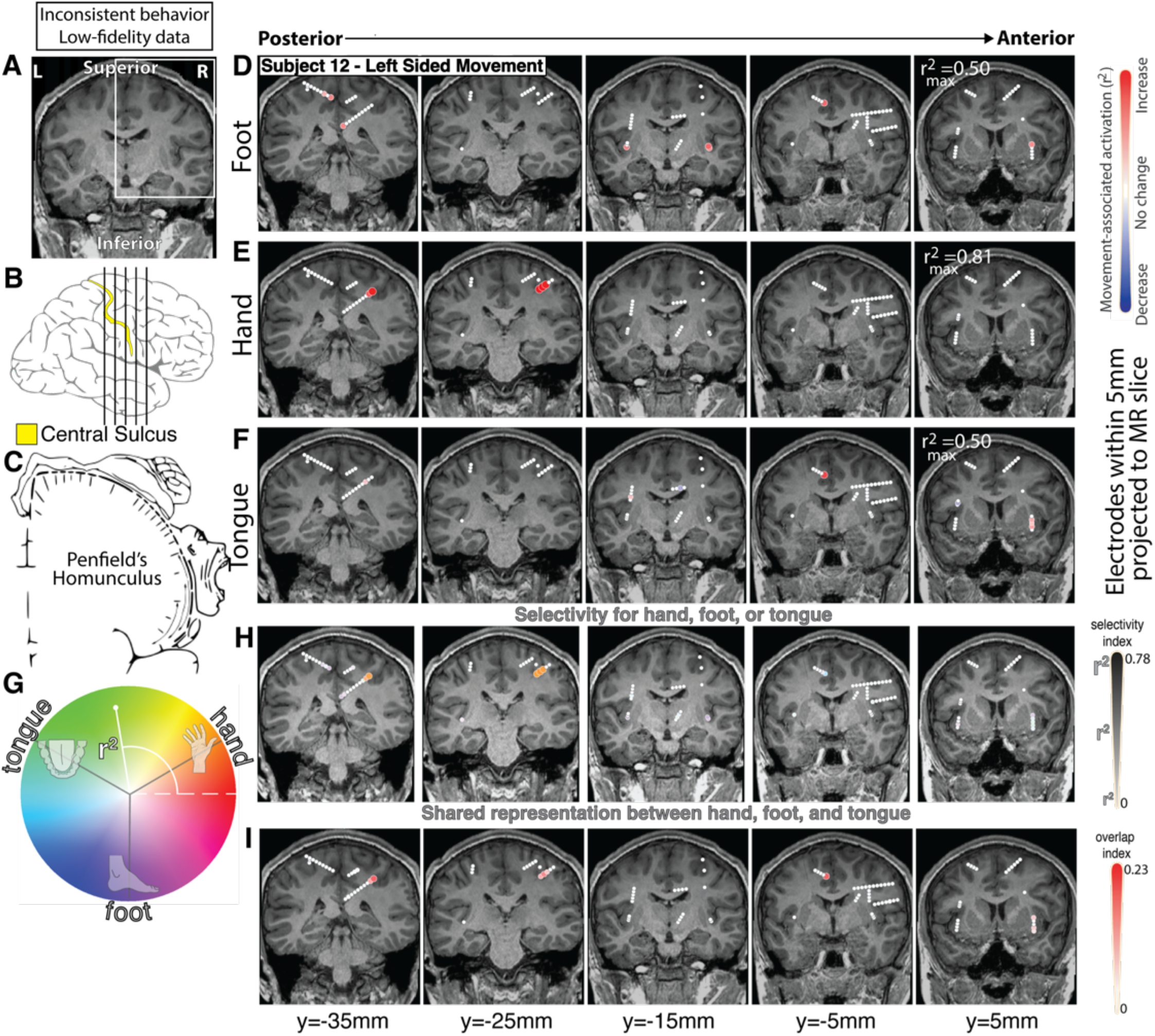
As in the main text Figure 2, for subject 12 during <u>left-sided</u> movement.

**Figure S27.**
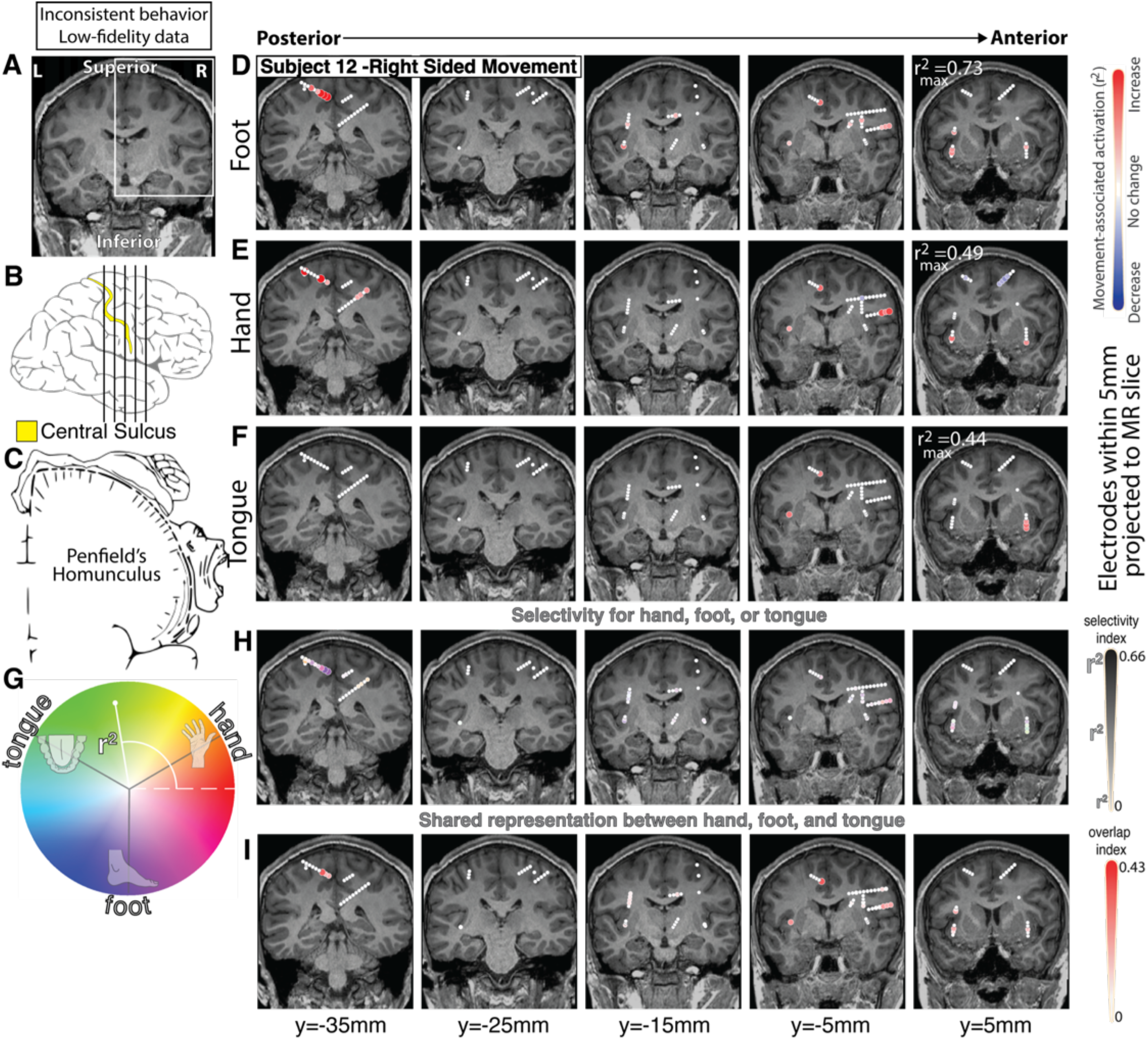
As in the main text Figure 2, for subject 12 during <u>right-sided</u> movement.

**Figure S28.**
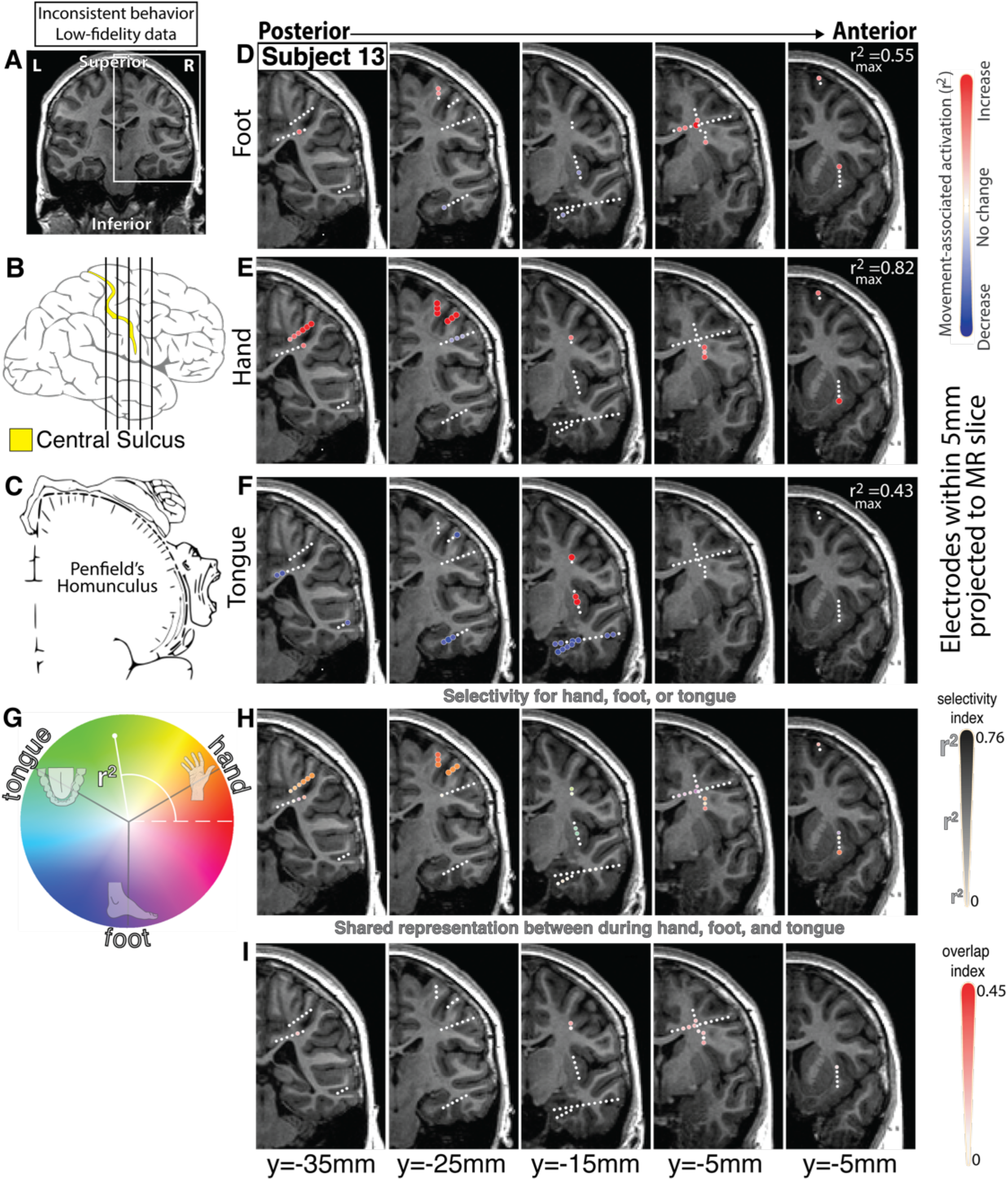
As in the main text Figure 2, for subject 13.

**Figure S29.**
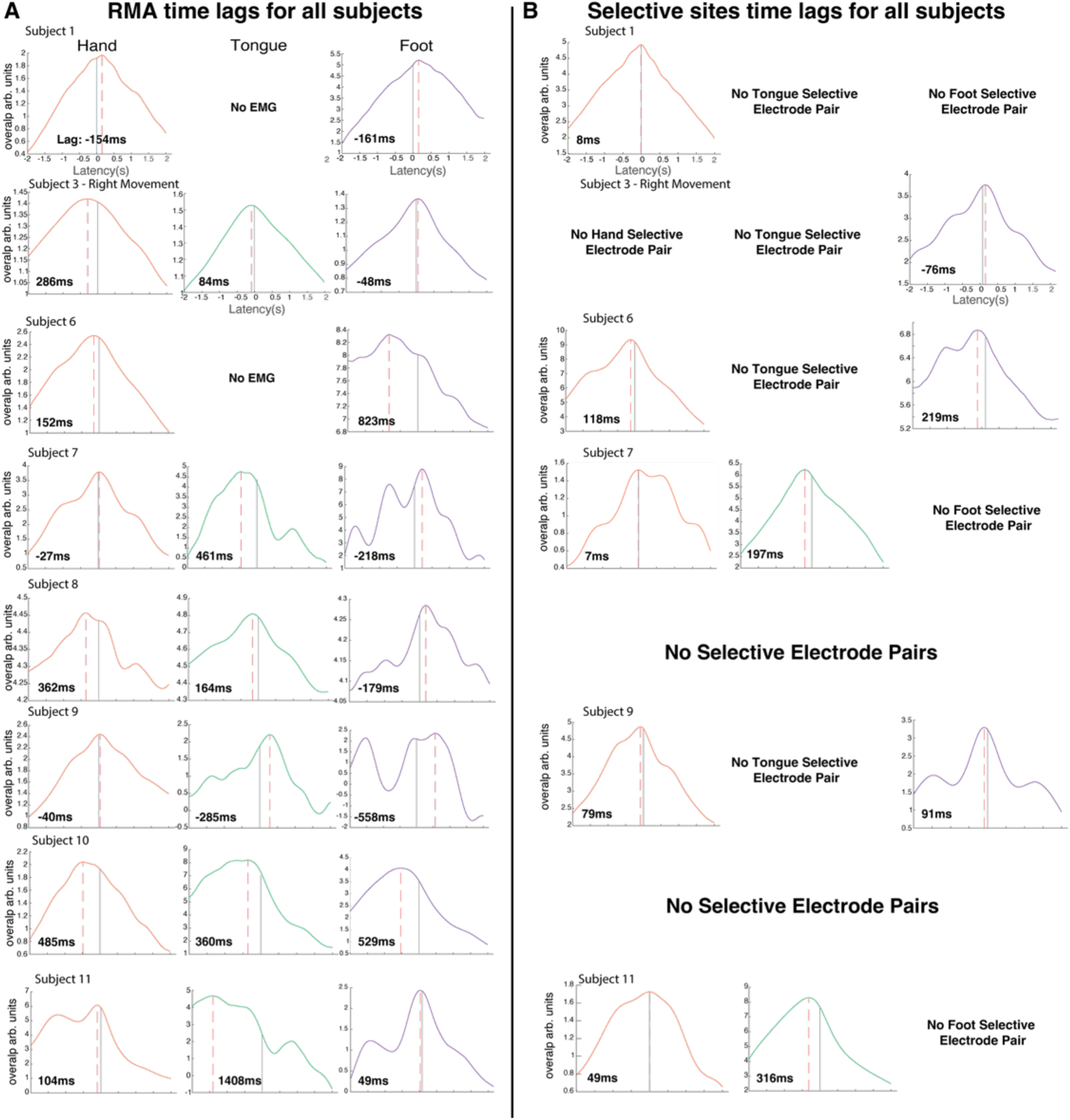
Latency plots for all subjects that have not already been shown in other figures. **A.** Time lag plots for all subjects between the RMA site and EMG for hand, tongue, and foot (labeled and colored). **B.** Time lag plots for all subjects between sites selective for certain modalities and EMG for hand, tongue, and foot (labeled and colored). Lag values are seen in the bottom left corner of each plot, with positive values representing broadband signals that occur prior to EMG.

**Table S2.**
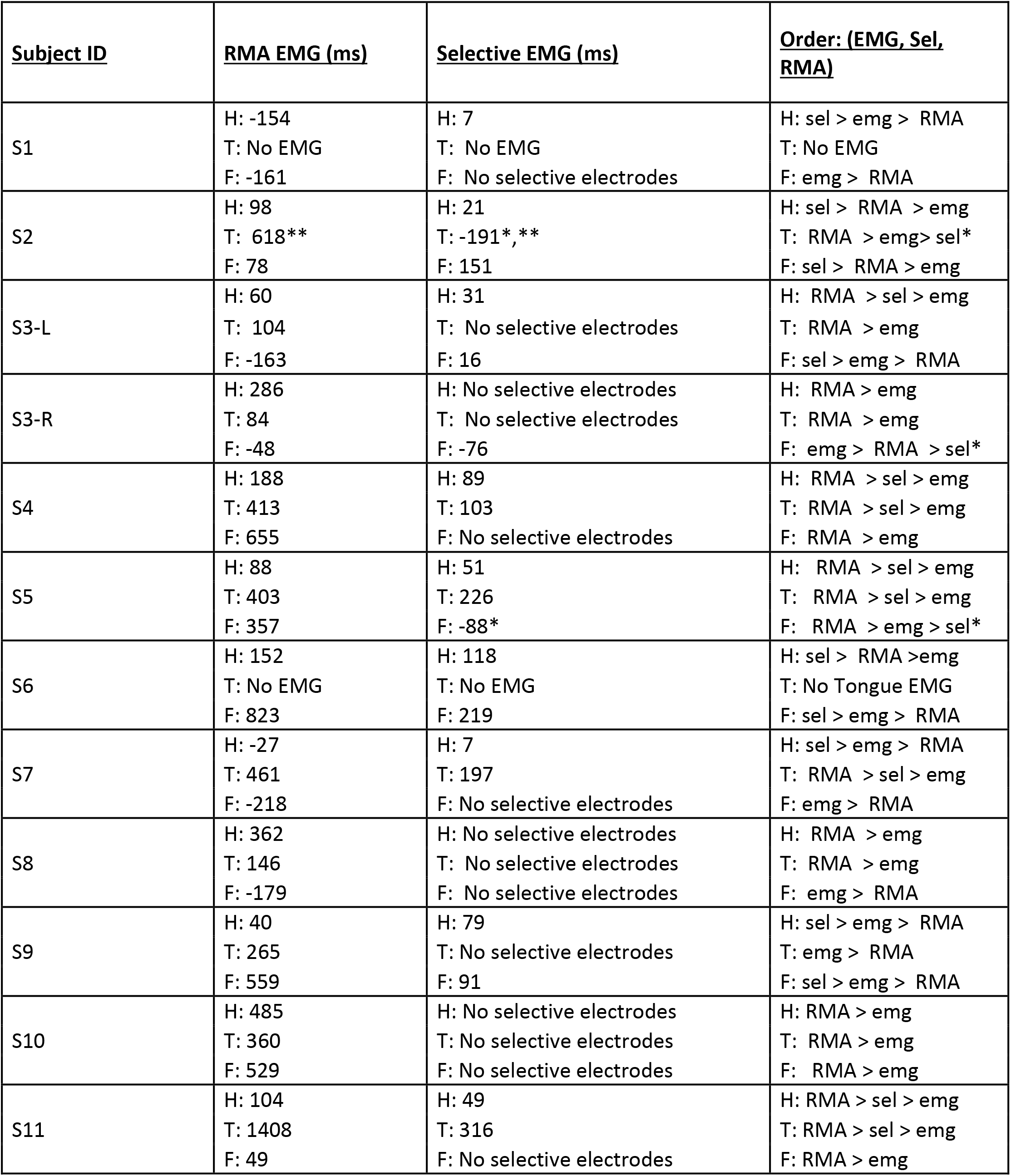
Timing of EMG, RMA, and Selective sEEG signals. Positive values indicate that sEEG signal peaks prior to EMG. * Most selective channel likely in sensory cortex ** Low EMG quality for initial trials

## Notes

https://osf.io/p5n2k

### Competing Interest Statement

The authors have declared no competing interest.

### Summary of Updates

We revised the initial upload to include the edge of 3 panels (H-J) that were previously cut off in Figure 4 of the main text.

